# Frustration in protein complexes leads to interaction versatility

**DOI:** 10.1101/2020.11.11.378091

**Authors:** Maria I. Freiberger, Peter G. Wolynes, Diego U. Ferreiro, Monika Fuxreiter

## Abstract

Disordered proteins can fold into a well-defined structure upon binding but these complexes are often fuzzy: the originally disordered partner adopts different binding modes when bound to different partners. Here we perform a systematic analysis of 160 proteins that form fuzzy complexes and demonstrate that the disordered partner displays a high degree of frustration in both the free and bound states. Although the folding of disordered regions upon binding reduces frustration relative to that of the unbound state, the interactions at the binding interface do not become fully optimized. In addition, we show that sub-optimal interactions lead to alternative frustration patterns in the complexes with different partners. These results demonstrate that disordered proteins do not always achieve fully optimal interactions in their complexes and their residual frustration leads to interaction versatility with different partners.

## Introduction

The discovery of disordered proteins has challenged the highly successful structure-function paradigm of molecular biology by raising the question of how biomolecular recognition can be achieved without a specific well-defined tertiary structure [24] [2]. Disordered proteins often function as interaction hubs in which the binding of multiple partners controls the specificity of signaling pathways [30]. While in the past two decades a series of experimental and computational methods have begun to characterise the conformational ensembles of disordered proteins in their free states [14], their properties in the bound state are less well understood. Disordered proteins often display different structures when they are bound with different partners. This phenomenon is termed “fuzzy binding” [8]. The observed binding modes of disordered proteins range from becoming nearly fully ordered to forming rather disordered states in the bound complex. The structures can also change through posttranslational modification or by varying cellular conditions [19]. Fuzzy binding enables disordered proteins to interact not with every biomolecule but specifically only with a defined set of partners. The physical basis of this controlled promiscuity has not yet been revealed.

Disordered proteins occupy a broad range of their energy landscapes. It has been established that conformational ensembles of disordered proteins are however not fully random. They rather form secondary structure elements [25], with many alternative intramolecular interactions leading to numerous but somewhat structurally distinct conformational sub-states in the native ensemble (Fig. 1). It is the entropic penalty of folding, which is a bottleneck for folding of disordered proteins.

**Figure 1.**
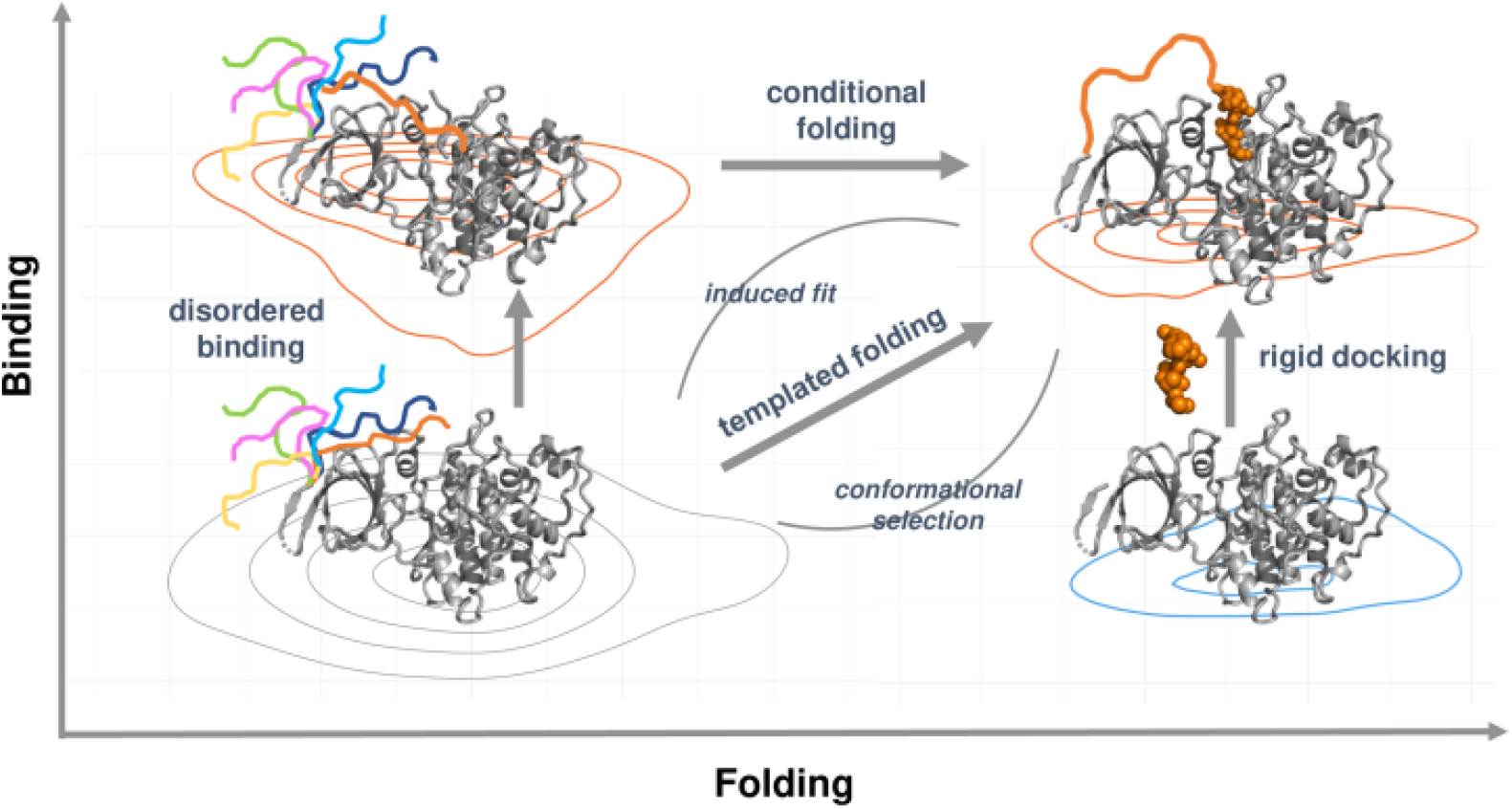
Schematic representation for folding and binding landscapes. When local frustration is low, the folded proteins associate as rigid bodies. When local frustration is high, folding is coupled to binding, templating the folding of the disordered region. If local frustration remains in the bound state, many conformations are still accessible leading to fuzziness. The structures illustrate different binding modes derived from structures of glycogen-synthase kinase 3 (GSK-3). The right side (vertical) represents rigid docking, when a folded protein binds to a folded partner, which in this case is the LRP6 peptide (orange) (PDB: 4nm5). The left side (vertical) represents disordered binding, when the disordered N-terminal region of GSK3 makes transient interactions with the active site. The different conformations are schematically represented by colored lines. The process displayed on top (from left to right) is the conditional folding, when the N-terminal peptide folds upon phosphorylation and binds the active site with a well-defined conformation. This structure (PDB: 4nm3) superimposes well on the complex with the LRP6 peptide (PDB: 4nm5). The orange line (top right) emphasizes that a part of the N terminal region remains to be disordered in the complex. The diagonal (from bottom left to top right) represents the templated folding, when the disordered region adopts a well-defined structure upon binding, which can be achieved via conformational selection or induced fit. This scenario is different from conditional folding when both ordered and disordered bound states can be observed.

Disordered proteins, can undergo templated folding upon binding with their partners, leading to a more well-defined conformation in the bound state [32]. Templated folding can be described by a funnel-like free energy landscape, which is made up from both intra- and intermolecular interactions, in contrast to autonomously folding proteins, whose funnel can be generated by intramolecular interactions alone (Fig. 1). Templated folding can sometimes be described as conformational selection [29], when one of the conformational sub-states already dominant in the free state is stabilized or may be termed induced fit, when the new conformation promoted by the partner is present only in very low concentration in the free ensemble [11]. According to either description the intermolecular interactions of disordered proteins with their partners are thought to contrast with the rugged landscape that would arise from their intramolecular interactions alone.

Templated folding differs from autonomous folding in that often a considerable portion of the protein remains disordered in the final bound complex [1] [10]. Thus, templated folding cannot always be described by a perfectly funnel-like energy landscape. Although the structural motifs found in the bound complex often overlap with the pre-formed conformational elements in the unbound state [9], distinct conformations turn out to be sampled when the protein binds to different partners. Furthermore, mutations, which stabilize binding competent secondary structure elements in the bound form may not always improve binding affinity [23]. Surprisingly, in these cases mutations outside the binding region often contribute to the affinity or specificity of binding [4]. These results suggest that heterogeneous nucleation in the templated folding pathway differs somewhat from the homogeneous nucleation of single globular proteins [28] [15].These results also suggest that the energy landscape of the bound complex is more rugged than for more fully folded species.

Disordered proteins can also undergo disordered binding, displaying many conformational sub-states in the bound forms, generating a rugged energy landscape. Conformational exchange between these sub-states can be observed both within and outside the binding region, [8] [20]. This pattern facilitates transient interactions at the binding interface with other functional motifs [22]. All these observations prompt the idea that the interactions of disordered proteins can be fuzzy, and that their functional versatility exploits the diversity of many different sub-states [27]. Although the biological significance of fuzziness has been established, understanding how diversity and specificity are reconciled requires the quantitative application of energy landscape theory. One’s intuition that interactions mediated by disordered regions must be always weak is contradicted by the existence of disordered protein complexes with high affinities [2].

Here we examine this problem using energy landscape theory tools that analyze local frustration in proteins [5]. These approaches were originally developed to describe how individual parts of a protein guide the folding of globular proteins towards their minimally frustrated native state [3]. Localizing frustration has given insights into their conformational motions and into their functional adaptations that conflict with folding [6]. In this paper we expand the theory of frustration to complexes of disordered proteins, showing consistency with the energy landscape theory. In this paper we systematically analyze frustration in the free and bound states of 160 disordered proteins that have been found to form fuzzy complexes. We find that the interactions display a high degree of frustration in both the more structured and unfolded parts of disordered proteins. We also show that while templated folding reduces the level of frustration, it does not eliminate frustration entirely, reflecting the fact that intermolecular interactions in the distinct fuzzy protein complexes are sub-optimal. We find there are often distinct frustration patterns in complexes with different partners, which indicates that using sub-optimal interactions provides some selectivity but also enables versatility. Our results provide a consistent physical model by which energetic frustration explains the functional versatility of fuzzy protein complexes on the basis of the energy landscape theory.

## Materials and Methods

The protocols underlying the datasets have been published, as well as the datasets themselves. Therefore here we only provide a brief description of the methods, which are detailed in previous works [12, 18].

### Regions representing templated folding (disorder-to-order binding mode, DORs, Table S1)

We collected crystal structures from the PDB that have resolution higher than 3 *A*, but that have missing electron density for at least 5 residues. We excluded protein sequences with post-translational modifications or that contained non-standard amino acids. We then collected all available bound-state structures, and filtered those, to find the disordered region in all the complexes. We analyzed the interface residues, and selected those regions, where at least 1 residue mediated an inter-protein interaction (within 4.5 *A* from the interface). The resulting disorder to order, DOR, dataset contained 97 non-redundant disordered regions, which were represented in 331 complexes, where the disordered regions folded upon binding (Table S1).

### Regions representing context-dependent binding modes (CDRs, Table S2)

Disordered regions that were observed to be as both folded and disordered in different complex structures with distinct binding partners were defined as context-dependent regions, CDR. We assembled context-dependent regions with a minimum length of 5 residues in the CDR dataset. This dataset contained 93 non-redundant disordered regions, represented in both ordered and disordered forms in 750 complex structures (1505 chains) (Table S1).

### Dataset of Fuzzy Protein Structures and Local Frustration

The protein structures for DOR and CDR class were downloaded from the Protein Data Bank (PDB, https://www.rcsb.org/) and the frustration patterns were calculated using the protein frustratometer software [21], [13], http://www.frustratometer.tk/).

## Results

We calculate the local frustration patterns using the Frustratometer server [21] which measures the contribution of specific interactions to the creation of a funneled folding landscape. To quantify how the local frustration patterns correspond with fuzziness we calculated the relative pair distribution functions *g*(*r*) for the locations of various classes of contacts classified by their frustration level with respect to the C*α* locations of the residues found in fuzzy regions. This allows us to see the connection of frustration to fuzziness.

### Templated folding leaves highly frustrated interactions in fuzzy protein complexes

We have evaluated the local frustration patterns in a set of 83 structures (97 fuzzy regions) of disordered protein complexes, where the disordered regions manifest disorder-to-order transitions on binding. Fig. 2A shows an example of the local frustration patterns in a fuzzy protein complex. The pair distribution functions for residues classified by configurational frustration index are displayed in Fig. 2B. The corresponding distributions for residues as classified by mutational frustration index are shown in Fig.S1 for all the complexes generated by templated folding. The distributions also include disordered regions which fold, but which do not mediate interactions with the partner. We found that protein regions that were originally disordered but that now adopt a well-defined structure upon binding still exhibit highly frustrated interactions between 2 and 4 A. The density of minimally frustrated contacts is also lower in these bound but fuzzy regions (Fig. 2B (Left)). These results indicate that the folding of disordered regions upon binding often remains far from being optimal. We have also found that those disorder-to-order regions (DORs) which have taken on order by templated folding, also display an enrichment in highly frustrated interactions with respect to the rest of the molecule (Fig. S1A). The interactions found in the structured regions of the same proteins (chosen as random controls, Methods) also show an enrichment of highly frustrated contacts which are slightly less frustrated than those of disordered regions (Fig. 2B (Right)). The frustration of the interactions of the ordered regions remains significantly higher than what is usually found in complexes formed from fully structured proteins [5]. These results indicate that templated folding of fuzzy regions also imposes constraints on the folded part of the protein.

**Figure 2.**
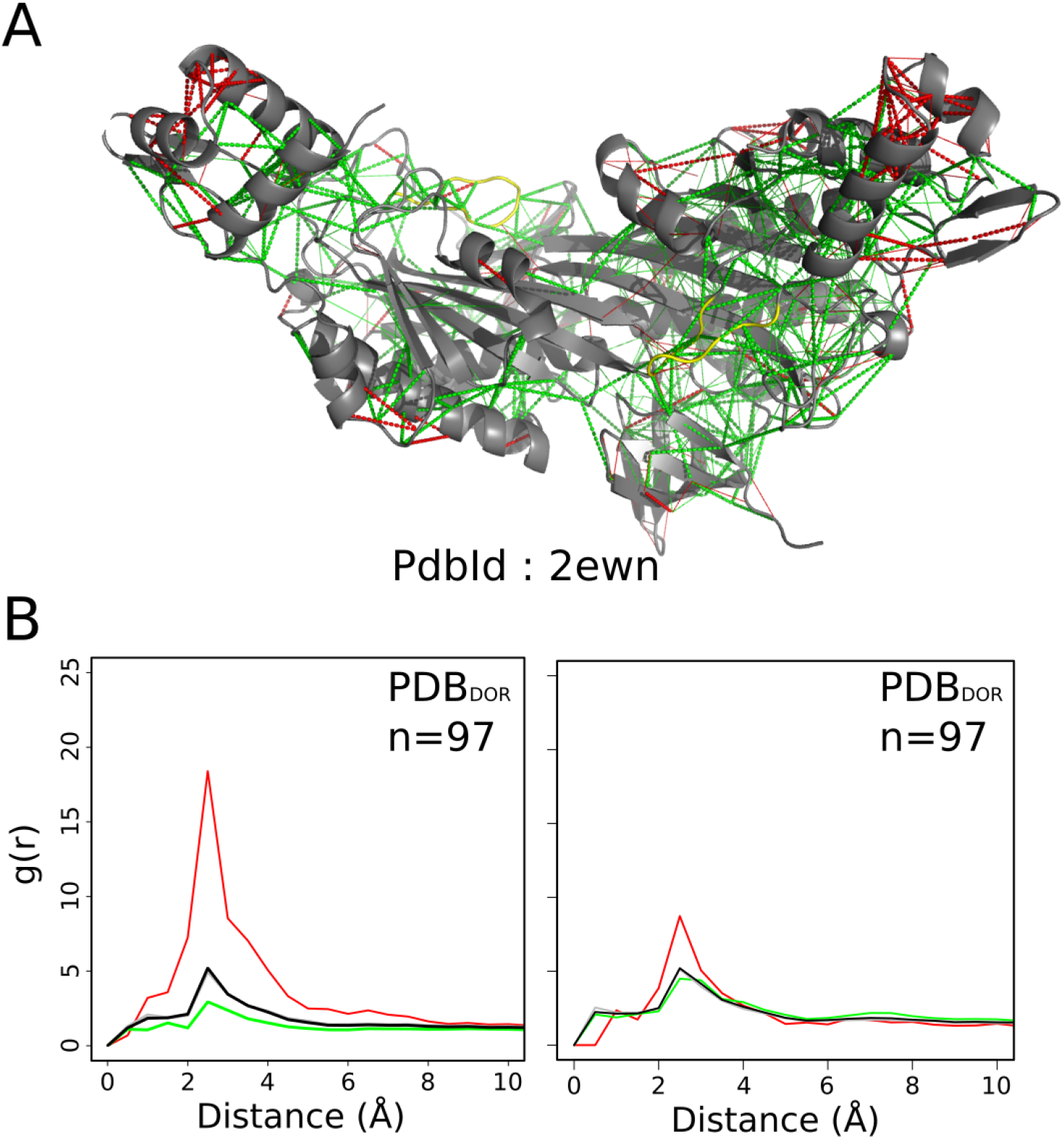
Local frustration in complexes of disordered proteins generated by templated folding where the regions undergoes transition from a disordered to an ordered state. A) Examples of frustration patterns in a protein undergoing disorder-to-order transition upon binding. The backbones of the protein are shown as gray cartoons, minimally frustrated contacts are depicted with green lines, highly frustrated interactions with red lines. Neutral interactions were omitted for clarity. The disorder-to-order region is colored yellow. B) On the left we show the pair distribution function of the contacts between the protein and the residues in the disorder-to-order region. On the right we show the pair distribution function of the contacts between residues of structured regions. Green: minimally frustrated contacts, red: highly frustrated, gray: neutral contacts, black: all contacts. In all cases g(r) values were normalized such that g(20)=1.

We next compared the level of frustration of those residues which are involved in the binding interface (binding) to those, which do not mediate intermolecular interactions (non-binding). Fig. 3A compares the density of the configurational frustration index for fuzzy residues involved in binding (blue) and non-binding contacts (pink), in the complexes generated by templated folding. We observed that those residues which do not form contacts with the partner exhibit higher frustration index than do those which directly form intermolecular contacts (Fig. 3A) indicating that the binding itself does ameliorate the high frustration of disordered proteins. These results indicate that folding of disordered regions is less optimal than their frustrated interface interactions.

**Figure 3.**
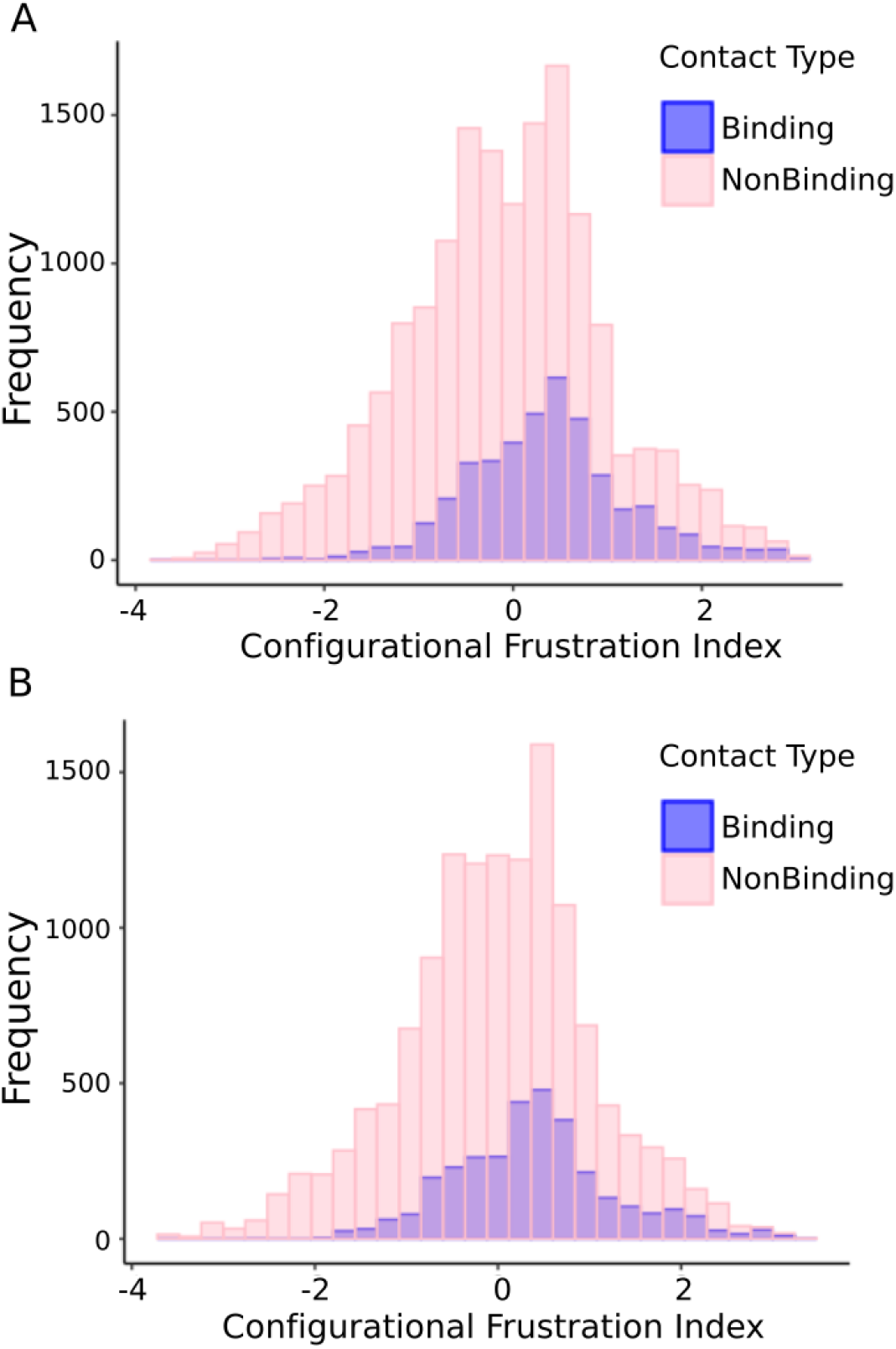
The distribution of local frustration indeces for binding contacts (red) and for the non-binding contacts (blue) for fuzzy residues. These are shown for A) Regions representing templated folding (DOR), and for B) Regions representing context-dependent binding modes (CDR).

### Templated folding decreases the overall frustration of disordered regions relative to the free monomeric state

Without simulating the intrinsically disordered protein ensemble it is difficult to assess precisely the local frustration of the disordered protein regions in their free (unbound) forms. We can get an idea of the frustration level however by examining the frustration in the structure of the disordered monomers simply by removing the interaction partner, thus generating a hypothetical single structure representative of the ensemble without intermolecular interactions. Both the finally disordered and structured regions display a higher density of frustrated contacts in the absence of the partner (Fig. S2). When we compare the frustration in monomers and complexes we see that the level of frustration is lowered upon binding: partner interactions do reduce the number of sub-optimal contacts.

For templated folding (Fig S3) and context-dependent binding modes (Fig S4) we observe that fuzzy binding also reduces frustration as compared with the free state. These results suggest that highly frustrated interactions are related to changes in binding modes.

We also analyzed some monomers that are found to be ordered in the free form but that become fuzzy in the bound form (Fig. S5). For these 52 regions we also observe an enrichment of highly frustrated interactions around the fuzzy regions.

### Conditional folding increases frustration of disordered regions

Increasing experimental evidence indicates that the frustrated interactions of disordered proteins [18] often manifest themselves by forming ordered complexes with some partners but forming disordered complexes with other partners [26]. We term these examples as displaying “conditional folding”. In this scenario, the folding of disordered proteins depends on the binding context (context-dependent regions, CDRs), such as the interaction partner, posttranslational modification or cellular conditions [16]. Here we analyzed 77 complexes (93 fuzzy regions), generated by conditional folding and found that these complexes exhibit more highly frustrated interactions (Fig. 4A) than the complexes generated by templated folding (Figure 2B). The distributions for the mutational frustration index are shown in Fig.S1. These indicate a small enrichment in frustrated interactions around the fuzzy regions. Frustrated contacts can be found both in the structured regions of the proteins (Figure 4B), and in the fuzzy regions outside the binding interface (Figure 3B). Thus, varying the degrees of folding with different partners results in sub-optimal interactions in the bound state.

**Figure 4.**
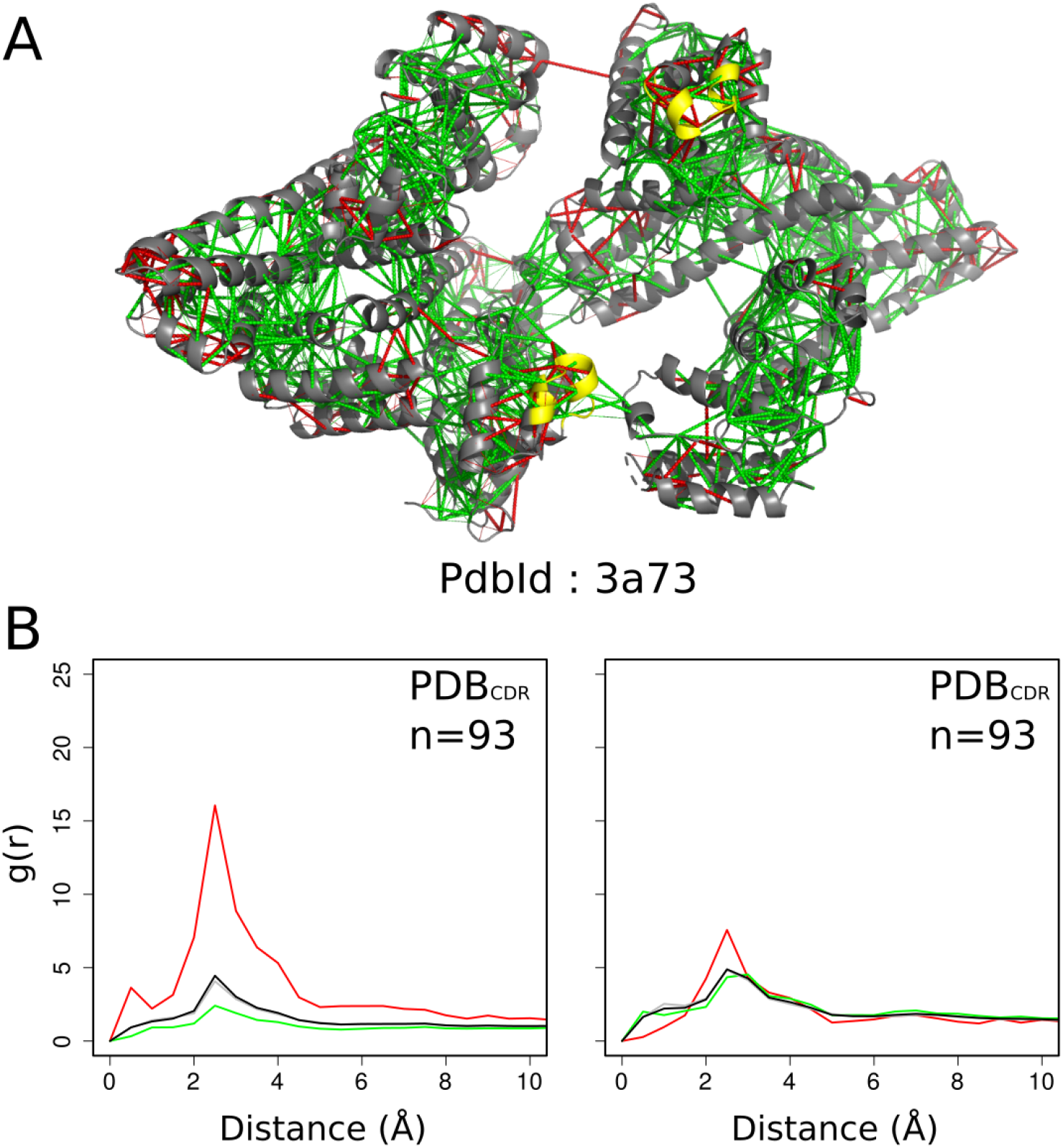
Local frustration in conditionally folding proteins. Those proteins which do not have a well-defined structure as monomer, but that may adopt a structure in a partner-or context-dependent manner or remain disordered in their complexes. A) Examples of frustration patterns in a conditionally folding protein. The backbone of the protein is shown as gray cartoons, minimally frustrated contacts are depicted with green lines, highly frustrated interactions with red lines. Neutral interactions were omitted for clarity. The context-dependent region is marked in yellow. B) On the left the pair distribution function of the contacts between the protein and residues of the context-dependent regions. On the right the pair distribution function of the contacts between residues of structured regions. Green: minimally frustrated contacts, red: highly frustrated, gray: neutral contacts, black: all contacts. g(r) values were normalized such that g(20).

### Partner-specific frustration facilitates target selection

The above results are consistent with the idea that the folding of disordered regions upon their targets does not always result in complex structures that are completely optimal and free from conflicts. What is the molecular basis of target selection in the absence of a distinguished bound-state conformation? To answer this question, we examined some complexes in which the same disordered region interacts with different binding partners.

We illustrate the differential binding of fuzzy regions by residues 39 - 47 of Translation initiation factor 2 subunit gamma (Q980A5). In protein structures PDB: 3cw2 and PDB: 3i1f eif2g folds into two different conformations. While the structure of the fuzzy region in 3cw2 is stabilised by the intramolecular interactions between Glu39-Thr46 and Glu40-Gly44, the 3i1f structure is instead stabilised by a charge-charge interaction between Glu39-Arg43. In the 3ilf structure, the Gly44 main chain forms a hydrogen bond with Lys42 side-chain while in the 3cw2 structure Lys42 interacts with Asp-283 of the structured domain.

Fig 5A shows the structures and local frustration frustration patterns for these different protein structures. Overall both possess an extended network of minimally frustrated interactions, with patches of highly frustrated interactions on the surface. The structures each display different frustration patterns for the fuzzy region in the complexes. We see that different ways of resolving the conflicts have been chosen in the alternative structures. This interchange of frustrated interactions is easily visualized in the contact maps.

**Figure 5.**
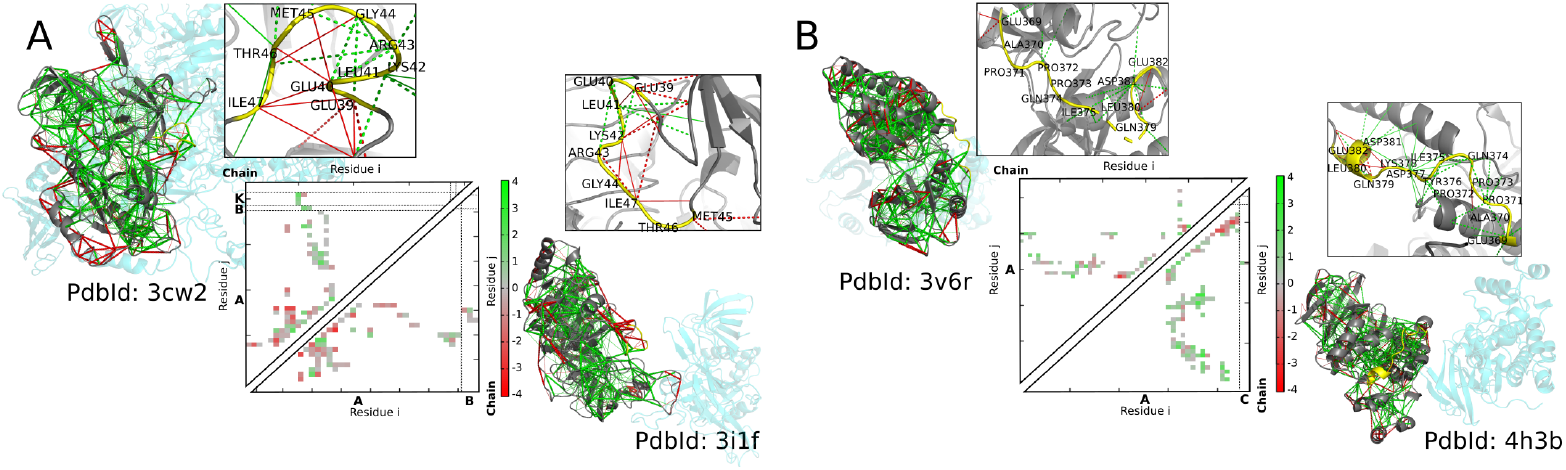
A) Structures for Translation initiation factor 2 subunit gamma (eif2g), PdbID: 3cw2 (above), and PdbID: 3i1f (below). Contact map of 3cw2 (above the diagonal), and 3i1f (below the diagonal). B) Protein structure for Mitogen-activated protein kinase 10, PdbID: 3v6r (above), and PdbID: 4h3b (below). Contact map of 3v6r (above the diagonal), and 4h3b (below the diagonal). The local frustration patterns of the protein, with the minimally frustrated interactions in green, the neutral in gray and highly frustrated interactions in red. The fuzzy region is shown with yellow backbone. For contact map, green: minimally frustrated contacts, red: highly frustrated, gray: neutral contacts. We see that which contacts are frustrated change in the alternate structures.

Another example (Fig 5B), that illustrates the nature of fuzzy binding is the 369 - 382 residue region of Mitogen-activated protein kinase 10 (P53779). In the PDB: 4h3b structure, the folding of the fuzzy region is stabilised by many intramolecular interactions. As we can observe in the contact map, some of these interactions form between side-chains with main-chain atoms, e.g. Gln374 and Pro372, Gln379 and Leu380, or Glu382 NE2 and Glu382 C-O. In contrast, in the PDB: 3v6r structure, considerably fewer intarmolecular contacts (Glu369, Leu380 and Asp381). The contact maps clearly reflect the different ways of trying to eliminate frustration in the two complexes.

## Discussion

Some frustration in the energy landscapes is required for the functional adaptability of proteins [6]. As a consequence of those frustrated interactions, the energy landscape is rugged encompassing distinct local minima enabling multiple biological activities [7]. To control their interactions generally proteins have evolved to optimize frustration allowing specificities to be compromised for functional needs [17]. This is brought to the extreme for disordered proteins, which even lack a well-defined conformation on their own and must always be described as an ensemble of conformers [30]. The structured character of complexes of disordered proteins with their specific partners however, has lead to the misleading impression that, in the end, functioning always requires a single well-defined conformation to be dominant. The presence of a well defined structure, however, does not correlate with the affinity of the interactions.

Here we have performed a systematic analysis of the complexes of many disordered proteins. This analysis demonstrates that even after binding, the interaction energetics are far from optimal in the fuzzy regions. Consistent with earlier results for individual cases [28], we have found that both the disordered and structured regions of complexes are enriched in highly frustrated interactions in the bound complexes of disordered proteins. The interface contacts do decrease the frustration level in the disordered protein once bound as compared to the frustration of the free state, but the interactions often still remain sub-optimal and energetic conflicts remain to be resolved. These results corroborate the ruggedness of the energy landscape, which describes complexes of disordered regions. We demonstrate that the fuzzy regions display distinct frustration patterns with different partners, rationalizing how residual frustration allows both specificity and versatility to be encoded. These observations can be exploited upon targeting disordered regions by small molecules. Our work shows that specificity of the interaction is not solely encoded in a given contact pattern, but also by the way frustration is ameliorated. These observations highlight the importance of conflicting, sub-optimal interaction in drug design for disordered regions.

The coupled folding and binding of disordered regions leads to sub-optimal contacts, which thus allows binding to different partners. This is fully consistent with the original notion of local frustration in spin glasses and systems like the triangular antiferromagnet where many structures compete as global minima [6] [31]. The high residual frustration explains why disordered regions are capable of manifesting several different binding modes [12]. The frustration of interactions in disordered proteins and their bound complexes allows binding to be fuzzy by being sub-optimal thereby enabling multifunctionality. Frustration and the ruggedness of the energy landscape thus enables functional versatility along with specificity.

## Acknowledgments

This work was supported by the Consejo de Investigaciones Cientificas y Tecnicas (CONICET), the Agencia Nacional de Promocion Cientifica y Tecnologica (Grant PICT2016/1467 to D.U.F.), and NASA Astrobiology Institute-Enigma (Grant Number: 80NSSC18M0093). D.U.F. is a CONICET researcher and M.I.F. holds a CONICET fellowship. Additional support was provided by the D. R. Bullard-Welch Chair at Rice University (Grant C-0016 to P.G.W.) M.F. acknowledges the financial support of HAS-11015, GINOP-2.3.2-15-2016-00044 and thanks to INFN Sezione di Padova for financial support.

