## Supplementary figures and tables for "Frustration in protein complexes leads to interaction versatility"

### References

### Supporting information

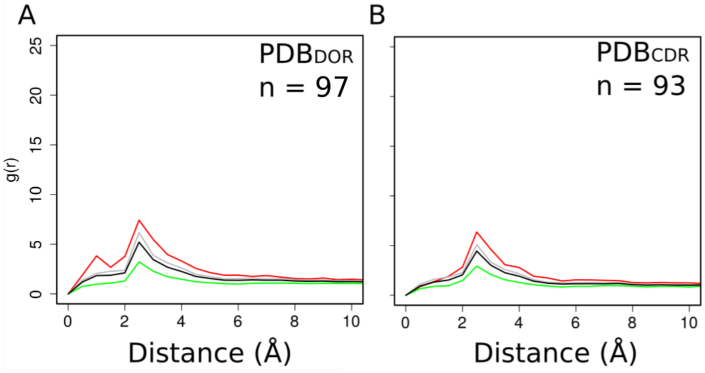


**Figure 1.** Mutational frustration of disordered regions in complexes generated by templated folding (A) and conditional folding (B). Pair distribution functions of the mutational frustration indices computed between the C$\alpha$ of the annotated fuzzy residue in the bound form. Green: minimally frustrated contacts, red: highly frustrated, gray: neutral contacts, black: all contacts. g(r) plots were adjusted in their axis ranges to enhance visualizations, however in all cases g(r) values were normalized such that g(20)=1.

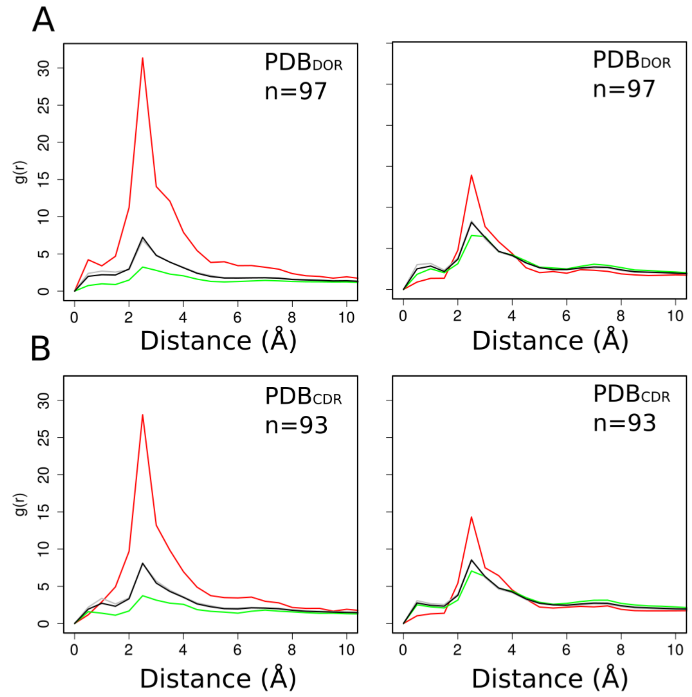


**Figure 2.** Configurational (left) and mutational (right) frustration of disorder-to-order regions (A) and context-dependent regions (B) in their unbound (monomeric) forms. (A) Disorder-to-order regions (DORs) undergo templated folding upon binding (B) Context-dependent regions conditionally fold upon binding to a specific set of partners. Pair distribution functions for single chains in the monomeric form were computed between the C$\alpha$'s of the annotated fuzzy residues and the contacts in different classes. The configurational frustration index is shown on the left and mutational frustration index is shown on the right. Green: minimally frustrated contacts, red: highly frustrated, gray: neutral contacts, black: all contacts. g(r) plots were adjusted in their axis ranges to enhance visualizations, however in all cases g(r) values were normalized such that g(20)=1.

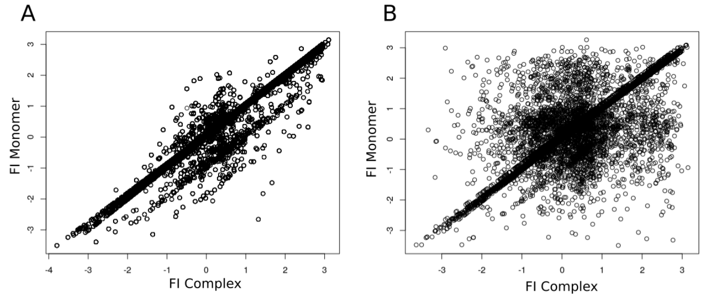


**Figure 3.** Templated folding reduces frustration of disordered regions as compared to their free forms. Correlation between the configurational frustration indices in the monomeric and bound forms are shown for contacts (A) involving residues of disorder-to-order (DOR) regions, (B) involving residues of structured regions.

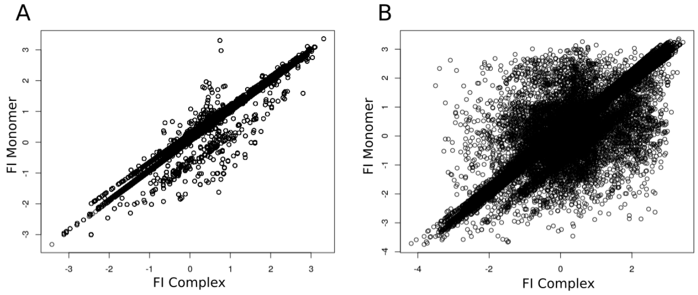


**Figure 4.** Conditional folding reduces frustration of disordered regions as compared to their free forms. Correlation between the configurational frustration indices in the monomeric and bound forms are shown for contacts (A) involving residues of context-dependent (CDR) regions, (B) involving residues of structured regions.

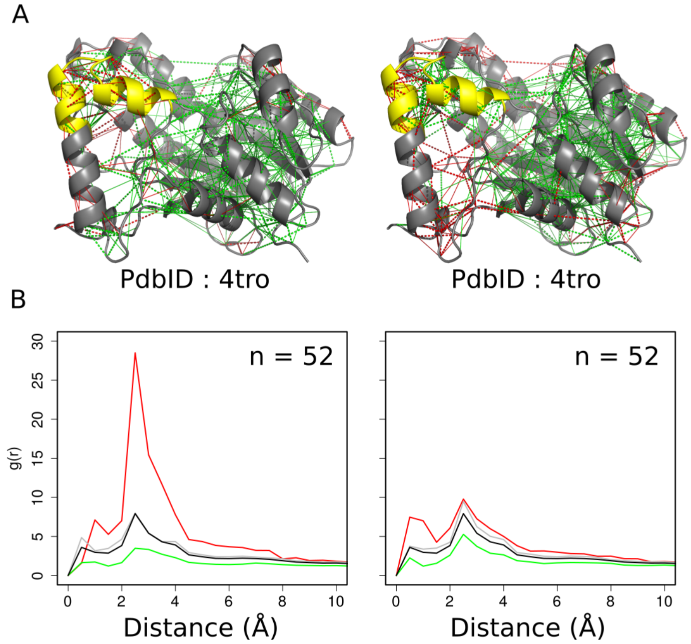


**Figure 5.** Frustration patterns of order-to-disorder regions. These regions are ordered in the unbound form and become disordered (unfold) upon binding. (A) Configurational (left) and mutational (right) frustration patterns are illustrated in the structure of enoyl-ACP reductase in complex with isoniazid (PDB:4tro). The backbones of the proteins are shown as gray cartoons, the fuzzy region is colored yellow (PBDID: 4tro, residues 197 - 215). The minimally frustrated contacts are depicted with green lines, highly frustrated interactions with red lines. Neutral interactions were omitted for clarity. (B) Pair distribution functions g(r) for configurational (left) and mutational (right) frustration indices, computed between the C$\alpha$ of the annotated fuzzy residue and the rest of the protein. Green: minimally frustrated contacts, red: highly frustrated, gray: neutral contacts, black: all contacts. g(r) plots were adjusted in their axis ranges to enhance visualizations, however in all cases g(r) values were normalized such that g(20)=1.

| UniprotID | PdbId | PdbId (g(r) analysis) | Fuzzy Residues (Uniprot) | Fuzzy Residues (Pdb) |
|---|---|---|---|---|
| A0QTN8 | 3DG7_B 3DG7_D | 3DG7_B | 17 - 27 | 17 - 27 |
| A5HZZ9 | 3DS9_A | 3DS9_A | 245 - 249 | 245 - 249 |
| B7TVP1 | 3GLM_A 3GLM_B 3GLM_C 3GLM_D | 3GLM_A | 237 - 250 | 237 - 250 |
| D0EM60 | 5F55_A 5F56_A | 5F55_A | 515 - 522 | 515 - 522 |
| D7PC21 | 3OU6_A 3OU6_B 3OU6_C 3OU6_D 3OU7_A 3OU7_B 3OU7_C 3OU7_D | 3OU6_A | 151 - 162 | 151 - 162 |
| F1RQI7 | 4E7Z_B | 4E7Z_B | 175 - 179 | 175 - 179 |
| I6Y9J2 | 4QRA_A 4QRA_B | 4QRA_A | 308 - 319 | 308 - 319 |
| O07347 | 1O87_A 1O87_B 1OKK_A 2CNW_A 2CNW_B 2CNW_C 2J7P_A 2J7P_B | 1O87_A | 272 - 277 | 272 - 277 |
| O14757 | 1NVQ_A 1NVR_A 1NVS_A 2YDJ_A | 1NVQ_A | 74 - 81 | 74 - 81 |
| O28126 | 1TFW_A 1TFW_B 1TFW_C 1TFW_D 2DR5_A 2DR7_A 2DR9_A 2DRA_A 2DRB_A 2DVI_A 2ZH1_A 2ZH2_A 2ZH3_A 2ZH5_A 2ZH6_A 2ZH8_A 2ZH9_A 3OUY_B 2ZH7_A | 1TFW_A | 89 - 95 | 89 - 95 |
| O43924 | 5TB5_B 5TB5_D 5E8F_A 5E8F_C 5F2U_A 5F2U_B 5TAR_B | 5TB5_B | 111 - 116 | 111 - 116 |
| O59282 | 3AEV_B | 3AEV_B | 26 - 32 | 26 - 32 |
| O60341 | 2Y48_A 5L3E_A 5LGT_A 2IW5_A 2UXN_A 2UXX_A 4XBF_A | 2Y48_A | 460 - 474 | 460 - 474 |
| O76074 | 2H42_B 1UDU_A 1UDU_B | 2H42_B | 791 - 807 | 791 - 807 |
| P00509 | 1ARG_A 1ARG_B 1ASL_A 1ASL_B 1ASM_A 1ASM_B 1ASN_A 1ASN_B 1X28_A 1X28_B 1X29_A 1X29_B 1X2A_A 1X2A_B 5VWQ_A | 1ARG_A | 10 - 17 | 10 - 17 |
| P00509 | $1ASM_B$ | $1ASM_B$ | 21 - 28 | 21 - 28 |
| P00592 | 1HN4_A 1HN4_B 1FX9_A 1FX9_B 1FXF_A 1FXF_B 1L8S_A 1L8S_B 3FVI_B 3FVI_C 3FVI_D | 1HN4_A | 38 - 43 | 38 - 43 |
| P00766 | 1T8L_A 1T8L_C 3T62_A 4Q2K_D | 1T8L_A | 71 - 76 | 71 - 76 |

Continues on next page.

| UniprotID | PdbId | PdbId (g(r) analysis) | Fuzzy Residues (Uniprot) | Fuzzy Residues (Pdb) |
|---|---|---|---|---|
| P00797 | 2IKO_A 2IKU_A 2IL2_A 2V0Z_C 2V16_C 3G72_B 3OOT_A 3OQF_A 3OQK_A 3Q3T_B 3Q4B_A 3Q4B_B 3Q5H_B 3SFC_A 3VSW_A 3VSX_A 3VUC_A 3VYD_A 3VYE_A 3VYF_A 4GJ5_B 4GJ6_B 4GJ7_A 4GJ8_B 4GJA_A 4GJB_B 4GJC_A 4GJD_A 4PYV_A 4Q1N_A 4RYC_A 4RYG_A 4RZ1_A 4S1G_A 4XX3_A 4XX4_A 5SY2_A 5SXN_A 5SY3_B 5SZ9_A 5T4S_A 1BIL_A 1BIM_A 1HRN_A 3GW5_B 3KM4_B 5KOQ_A 5KOS_A 5KOT_A 5TMG_A 5TMK_A 5V8V_B | 2IKO_A | 353 - 361 | 353 - 361 |
| P01112 | 1BKD_R 1K8R_A 1NVV_R 1NVW_Q 1NVW_R 1WQ1_R 1XD2_B 4NYJ_R | 1BKD_R | 66 - 71 | 66 - 71 |
| P02754 | 2R56_A 2R56_B | 2R56_A | 169 - 178 | 169 - 178 |
| P02945 | 1CWQ_A 1CWQ_B | 1CWQ_A | 244 - 252 | 244 - 252 |
| P02945 | 5VN7_B 5VN9_A 5VN9_B | 5VN7_B | 166 - 179 | 166 - 179 |
| P03252 | 1AVP_A 1NLN_A 4PID_A 4PIE_A 5FGY_A | 1AVP_A | 97 - 104 | 97 - 104 |
| P04547 | 1IXY_A 1IXY_B 1M5R_A 1M5R_B 1SXP_A 1SXP_B 1SXQ_A 1SXQ_B | 1IXY_A | 108 - 122 | 108 - 122 |
| P04547 | 1IXY_A 1IXY_B 1M5R_A 1M5R_B 1SXP_A 1SXP_B 1SXQ_A 1SXQ_B | 1IXY_A | 68 - 76 | 68 - 76 |
| P05326 | 1IPS_A 1IPS_B | 1IPS_A | 324 - 331 | 324 - 331 |
| P06129 | 1UJW_A | 1UJW_A | 594 - 601 | 594 - 601 |
| P06129 | 1UJW_A | 1UJW_A | 249 - 260 | 249 - 260 |
| P06129 | 1UJW_A | 1UJW_A | 77 - 82 | 77 - 82 |
| P06709 | 2EWN_A 2EWN_B | 2EWN_A | 116 - 124 | 116 - 124 |
| P07737 | 2PAV_P 2PBD_P 3CHW_P | 2PAV_P | 57 - 62 | 57 - 62 |
| P08174 | 1H03_P 1H03_Q | 1H03_P | 206 - 211 | 206 - 211 |
| P09527 | 1VG9_B 1VG9_D 1VG9_F 1VG9_H | 1VG9_B | 65 - 71 | 65 - 71 |

Continues on next page.

| UniprotID | PdbId | PdbId (g(r) analysis) | Fuzzy Residues (Uniprot) | Fuzzy Residues (Pdb) |
|---|---|---|---|---|
| P09651 | 1U1K_A 1U1L_A 1U1M_A 1U1N_A 1U1O_A 1U1P_A 1U1Q_A 1U1R_A 1PGZ_A 1PO6_A 2UP1_A | 1U1K_A | 94 - 99 | 94 - 99 |
| P0A386 | 2AXT_V 2AXT_v 5H2F_V 5H2F_v | 2AXT_V | 158 - 163 | 158 - 163 |
| P0A7G6 | 3CMV_C 3CMV_D 3CMV_F | 3CMV_C | 158 - 165 | 158 - 165 |
| P0A817 | 1P7L_A 1P7L_B 1P7L_C 1P7L_D 1RG9_A 1RG9_B 1RG9_C 1RG9_D | 1P7L_A | 103 - 108 | 103 - 108 |
| P10824 | 1GP2_A | 1GP2_A | 10 - 31 | 10 - 31 |
| P11838 | 1EPL_E 1EPM_E 1ER8_E 1GVU_A 1OEX_A 2ER0_E 2ER7_E 2ER9_E 3ER5_E 4ER4_E 4LP9_A | 1EPL_E | 166 - 174 | 166 - 174 |
| P11838 | 1EPL_E 1EPM_E 1ER8_E 1GVU_A 1OEX_A 2ER0_E 2ER7_E 2ER9_E 3ER5_E 4ER4_E 4LP9_A | 1EPL_E | 200 - 208 | 200 - 208 |
| P17169 | 2J6H_A 2J6H_B | 2J6H_A | 603 - 609 | 603 - 609 |
| P17291 | 1FJG_G 1N32_G | 1FJG_G | 147 - 151 | 147 - 151 |
| P19400 | 1F34_B | 1F34_B | 138 - 142 | 138 - 142 |
| P19400 | 1F34_B | 1F34_B | 153 - 157 | 153 - 157 |
| P19821 | 3KTQ_A 3LWL_A 3LWM_A 3M8R_A 3M8S_A 3OJS_A 3OJU_A 3RR7_A 3RR8_A 3RRG_A 3RRH_A 3RTV_A 3SV3_A 3SV4_A 3SYZ_A 3SZ2_A 3T3F_A 4C8K_A 4C8L_A 4C8M_A 4C8N_A 4C8O_A 4CCH_A 4DF4_A 4DF8_A 4DFJ_A 4DFK_A 4DFM_A 4DFP_A 4DLG_A 4ELT_A 4ELU_A 4ELV_A 1QTM_A 4KTQ_A 2KTQ_A | 3KTQ_A | 504 - 512 | 504 - 512 |
| P20058 | 1QJS_B | 1QJS_B | 124 - 130 | 124 - 130 |
| P20783 | 3BUK_A 3BUK_B | 3BUK_A | 143 - 148 | 143 - 148 |
| P20783 | 3BUK_A 3BUK_B | 3BUK_A | 180 - 185 | 180 - 185 |
| P20783 | 3BUK_A 3BUK_B | 3BUK_A | 198 - 203 | 198 - 203 |
| P22301 | 1J7V_L 1Y6K_L | 1J7V_L | 56 - 62 | 56 - 62 |
| P23470 | 5E5R_A 5E5R_C | 5E5R_A | 292 - 297 | 292 - 297 |
| P23940 | 1BHM_A 1BHM_B 1ESG_A 1ESG_B 2BAM_A 2BAM_B 3BAM_A 3BAM_B | 1BHM_A | 79 - 91 | 79 - 91 |

Continues on next page.

| UniprotID | PdbId | PdbId (g(r) analysis) | Fuzzy Residues (Uniprot) | Fuzzy Residues (Pdb) |
|---|---|---|---|---|
| P24941 | 1F5Q_A 1F5Q_C 1FIN_A 1FIN_C 1FVV_A 1FVV_C 1OKV_A 1OKV_C 1OKW_A 1OKW_C 1OL1_A 1OL1_C 1OL2_A 1OL2_C 1URC_A 1URC_C 2C5N_A 2C5N_C 2C5O_A 2C5O_C 2C5V_A 2C5V_C 2C5X_A 2C5X_C 2I40_A 2I40_C 2UUE_A 2UUE_C 2V22_A 2V22_C 2WEV_A 2WEV_C 2WFY_A 2WFY_C 2WHB_A 2WHB_C 2X1N_A 2X1N_C 3F5X_A 3F5X_C 4FX3_A 4FX3_C 5IF1_A 5IF1_C | 1F5Q_A | 147 - 159 | 147 - 159 |
| P25685 | 3AGZ_A 3AGZ_B | 3AGZ_A | 336 - 340 | 336 - 340 |
| P28482 | 4XJ0_B | 4XJ0_B | 175 - 189 | 175 - 189 |
| P28631 | 1JR3_E | 1JR3_E | 259 - 264 | 259 - 264 |
| P29218 | 1IMA_A 1IMA_B 1IMB_A 1IMB_B 1IMC_A 1IMC_B 1IMD_A 1IMD_B 1IME_A 1IME_B 4AS4_A 4AS4_B 1AWB_A 1AWB_B 2HHM_A 2HHM_B | 1IMA_A | 31 - 39 | 31 - 39 |
| P31570 | 1G64_A 1G64_B 4HUT_A 4HUT_B | 1G64_A | 184 - 196 | 184 - 196 |
| P35555 | 1UZJ_B | 1UZJ_B | 1539 - 1548 | 1539 - 1548 |
| P36897 | 1B6C_B 1B6C_D 1B6C_F 1B6C_H 1IAS_A 1IAS_B 1IAS_C 1IAS_D 1IAS_E 3KCF_A 3KCF_B 3KCF_C 3KCF_D 3KCF_E | 1B6C_B | 175 - 199 | 175 - 199 |
| P40136 | 1K90_B | 1K90_B | 580 - 590 | 580 - 590 |
| P42700 | 2I91_A 2I91_B | 2I91_A | 336 - 340 | 336 - 340 |

Continues on next page.

| UniprotID | PdbId | PdbId (g(r) analysis) | Fuzzy Residues (Uniprot) | Fuzzy Residues (Pdb) |
|---|---|---|---|---|
| P49137 | 4TYH_A 2PZY_A 3A2C_A 3A2C_B 3A2C_C 3A2C_D 3A2C_E 3A2C_F 3A2C_G 3A2C_H 3A2C_I 3A2C_J 3A2C_K 3A2C_L 3KC3_A 3KC3_B 3KC3_C 3KC3_D 3KC3_E 3KC3_F 3KC3_G 3KC3_H 3KC3_I 3KC3_J 3KC3_K 3KC3_L 3WI6_A 3WI6_B 3WI6_C 3WI6_D 3WI6_E 3WI6_F 3R2B_A 3R2B_B 3R2B_C 3R2B_D 3R2B_G | 4TYH_A | 230 - 235 | 230 - 235 |
| P52732 | 1II6_B 2GM1_B 2GM1_D 2PG2_A 2PG2_B 2UYI_A 2UYI_B 2UYM_A 2UYM_B 2X2R_A 2X2R_B 2X2R_C 2XAE_A 2XAE_B 2XAE_C 3K5E_A 3K5E_B 3L9H_A 3L9H_B 4A51_C 4A51_D 4A51_G 4A5Y_A 4A5Y_C | 1II6_B | 174 - 181 | 174 - 181 |
| P53779 | 3V6R_B 3V6S_B 3OXI_A 4H36_A 4H39_A 4H3B_A 4H3B_C | 3V6R_B | 364 - 368 | 364 - 368 |
| P53779 | 4H3B_C | 4H3B_C | 71 - 75 | 71 - 75 |
| P71707 | 5CRF_A 5CRF_B 5CRF_C 5CRF_D | 5CRF_A | 780 - 785 | 780 - 785 |
| P74873 | 1G4U_S | 1G4U_S | 196 - 206 | 196 - 206 |
| P74873 | 1G4U_S | 1G4U_S | 220 - 224 | 220 - 224 |
| P74873 | 1G4U_S | 1G4U_S | 248 - 252 | 248 - 252 |
| P84022 | 1MK2_A | 1MK2_A | 324 - 328 | 324 - 328 |
| P84022 | 1MK2_A | 1MK2_A | 381 - 387 | 381 - 387 |
| P9WPN8 | 5LI6_A 5LI6_B 5LI7_A 5LI7_B 5LIE_A 5LIE_B | 5LI6_A | 75 - 79 | 75 - 79 |
| Q00987 | 4HBM_A 4HBM_B | 4HBM_A | 6 - 11 | 6 - 11 |
| Q07960 | 1OW3_A | 1OW3_A | 403 - 412 | 403 - 412 |
| Q09028 | 4PBY_A 4PBY_B 4PBZ_A 4PC0_A 4PC0_B | 4PBY_A | 16 - 23 | 16 - 23 |
| Q12341 | 4PSW_A | 4PSW_A | 200 - 208 | 200 - 208 |
| Q13325 | 4HOR_A 4HOT_A | 4HOR_A | 190 - 195 | 190 - 195 |
| Q14012 | 4FG9_A | 4FG9_A | 278 - 293 | 278 - 293 |
| Q14671 | 1M8W_A 1M8W_B 1M8X_A 1M8X_B 1M8Y_A 1M8Y_B 3Q0L_A 3Q0L_B 3Q0M_A 3Q0M_B 3Q0N_A 3Q0N_B 3Q0O_A 3Q0O_B 3Q0P_A 3Q0P_B | 1M8W_A | 1150 - 1165 | 1150 - 1165 |

Continues on next page.

| UniprotID | PdbId | PdbId (g(r) analysis) | Fuzzy Residues (Uniprot) | Fuzzy Residues (Pdb) |
|---|---|---|---|---|
| Q16539 | 5ETA_A 5ETA_B | 5ETA_A | 115 - 122 | 115 - 122 |
| Q45488 | 1D2I_A 1D2I_B 1DFM_A 1DFM_B | 1D2I_A | 115 - 122 | 115 - 122 |
| Q45488 | 1D2I_A 1D2I_B 1DFM_A 1DFM_B | 1D2I_A | 38 - 44 | 38 - 44 |
| Q60176 | 1HYG_A 1HYG_B | 1HYG_A | 224 - 229 | 224 - 229 |
| Q74P24 | 4EI7_A | 4EI7_A | 377 - 383 | 377 - 383 |
| Q74P24 | 4EI7_A | 4EI7_A | 80 - 88 | 80 - 88 |
| Q84AF2 | 2P0J_A 2P0J_B | 2P0J_A | 42 - 46 | 42 - 46 |
| Q8PJX9 | 4FOU_A 4FOU_B | 4FOU_A | 429 - 434 | 429 - 434 |
| Q8RNV8 | 2Q10_A 2Q10_B 3IMB_A 3IMB_B 3IMB_C 3IMB_D | 2Q10_A | 99 - 103 | 99 - 103 |
| Q8WTS6 | 1H3I_A 1H3I_B | 1H3I_A | 70 - 78 | 70 - 78 |
| Q96S59 | 5JIU_A 5JIU_B | 5JIU_A | 141 - 146 | 141 - 146 |
| Q9CCZ4 | 2CKD_B | 2CKD_B | 58 - 69 | 58 - 69 |
| Q9K2N0 | 5ACV_A 5ACV_B 5ACW_A 5ACW_B 5ACX_A 5ACX_B 5LM6_A 5LM6_B 5NHZ_A 5NHZ_B 5NI0_A 5NI0_E | 5ACV_A | 32 - 38 | 32 - 38 |
| Q9U1E1 | 3NGS_B 3NGT_A 3NGT_B 3NGT_C 3NGT_D 3NGT_E 3NGT_F 3NGT_G 3NGT_H 3NGT_I 3NGT_J 3NGT_K 3NGT_L | 3NGS_B | 145 - 150 | 145 - 150 |
| Q9X1H7 | 3PQC_A | 3PQC_A | 47 - 59 | 47 - 59 |
| Q9Y5B9 | 4Z2M_B | 4Z2M_B | 745 - 750 | 745 - 750 |

**Table 1.** Structures of complexes, which were generated by disorder-to-order transitions of protein regions (PDB$_DOR$). PDBIDs for all complexes are listed, those used for g(r) analysis are shown in a separate column. The sequence numbers of DOR regions are given based on both Uniprot and PDB numbering.

| UniprotID | PdbId | PdbId (g(r) analysis) | PDB complex, not structured | Fuzzy Residues (Uniprot) | Fuzzy Residues (Pdb) |
|---|---|---|---|---|---|
| A7ZS61 | 3GCM_A 3GCM_B 3GCM_C | 3GCM_A | 3GME_A | 275 - 296 | 275 - 296 |
| C8CHL4 | 3ZN5_A 3ZN5_B 3ZN5_C 3ZN5_D 3ZN5_E 3ZN5_F 3ZN5_G 3ZN5_H | 3ZN5_A | 3ZN6_B | 2 - 18 | 2 - 18 |
| O14757 | 1NVQ_A 1NVR_A 1NVS_A 2E9V_B 2YDJ_A | 1NVQ_A | 2YDJ_B | 2 - 8 | 2 - 8 |

Continues on next page.

| UniprotID | PdbId | PdbId (g(r) analysis) | PDB complex, not structured | Fuzzy Residues (Uniprot) | Fuzzy Residues (Pdb) |
|---|---|---|---|---|---|
| O14965 | 4C3P_A 4C3P_D | 4C3P_A | 3EFW_A 3EFW_B 3W18_A 3W18_B 3W2C_A 3W2C_C 3W2C_E 3W2C_G | 279 - 290 | 279 - 290 |
| O43189 | 5XFP_B | 5XFP_B | 5XFP_E | 322 - 326 | 322 - 326 |
| O50657 | 5GJM_A | 5GJM_A | 5GJM_B | 141 - 149 | 141 - 149 |
| O76074 | 1UDU_A 1UDU_B 2H42_B | 1UDU_A | 2H42_C | 665 - 676 | 665 - 676 |
| O95831 | 4BUR_A 4BUR_B | 4BUR_A | 4BUR_C 4BUR_D | 551 - 558 | 551 - 558 |
| P00489 | 1PYG_A 1PYG_B 1PYG_C 1PYG_D 3E3N_A 3E3N_B 3E3N_D 3E3N_F 3E3N_G 3E3N_H | 1PYG_A | 3E3L_A 3E3L_B 3E3L_C 3E3L_D 3E3N_C 3E3N_E | 315 - 325 | 315 - 325 |
| P00489 | 1PYG_A 1PYG_B 1PYG_C 1PYG_D 3E3N_A 3E3N_B 3E3N_D 3E3N_F 3E3N_G 3E3N_H | 1PYG_A | 3E3L_A 3E3L_B 3E3L_C 3E3L_D 3E3N_C 3E3N_E | 7 - 12 | 7 - 12 |
| P00492 | 1BZY_A 1BZY_B 1BZY_C 1BZY_D 4RAB_A 4RAB_B 4RAB_C 4RAB_D 4RAC_A 4RAC_B 4RAC_C 4RAC_D 4RAD_A 4RAD_B 4RAD_C 4RAD_D 4RAD_E 4RAD_F 4RAD_G 4RAD_H 4RAN_A 4RAN_B 4RAN_C 4RAN_D 4RAO_B 4RAO_C 4RAO_D 4RAQ_B 4RAQ_D | 1BZY_A | 1HMP_B 1Z7G_A 1Z7G_B 1Z7G_C 1Z7G_D 3GEP_A 3GEP_B 3GGC_A 3GGC_B 3GGJ_B 4RAO_A 4RAQ_A 4RAQ_C | 104 - 123 | 104 - 123 |
| P00514 | 3FHI_B | 3FHI_B | 3PNA_A 3PNA_B | 237 - 242 | 237 - 242 |
| P00514 | 3FHI_B | 3FHI_B | 3PNA_A 3PNA_B | 93 - 110 | 93 - 110 |

Continues on next page.

| UniprotID | PdbId | PdbId (g(r) analysis) | PDB complex, not structured | Fuzzy Residues (Uniprot) | Fuzzy Residues (Pdb) |
|---|---|---|---|---|---|
| P00766 | 1GL1_B 2CGA_B | 1GL1_B | 1ACB_E 1GL0_E 1GL1_A 1GL1_C 1OXG_A 1P2M_A 1P2M_C 1P2N_A 1P2N_C 1P2O_A 1P2O_C 1P2Q_A 1P2Q_C 1T7C_A 1T7C_C 1T8L_A 1T8L_C 1T8M_A 1T8M_C 1T8N_A 1T8N_C 1T8O_A 1T8O_C 2Y6T_A 2Y6T_B 2Y6T_C 2Y6T_D 3T62_A 3T62_B 3T62_C 4Q2K_A 4Q2K_B 4Q2K_C 4Q2K_D 5J4Q_A 5J4S_A | 11 - 15 | 11 - 15 |

Continues on next page.

| UniprotID | PdbId | PdbId (g(r) analysis) | PDB complex, not structured | Fuzzy Residues (Uniprot) | Fuzzy Residues (Pdb) |
|---|---|---|---|---|---|
| P00797 | 1BIL_A 1BIL_B 1BIM_A 1BIM_B 1HRN_A 1HRN_B 2IKO_A 2IKO_B 2IKU_A 2IKU_B 2IL2_A 2IL2_B 2V0Z_C 2V0Z_O 2V16_O 3G72_B 3OOT_A 3OOT_B 3OQF_A 3OQF_B 3OQK_A 3OQK_B 3SFC_A 3SFC_B 3VSW_A 3VSW_B 3VSX_A 3VSX_B 3VUC_A 3VUC_B 3VYD_A 3VYD_B 3VYE_A 3VYE_B 3VYF_B 4GJ7_A 4GJ7_B 4GJA_A 4GJA_B 4GJC_B 4GJD_A 4GJD_B 4PYV_B 4Q1N_A 4Q1N_B 4RYC_A 4RYC_B 4RYG_A 4RYG_B 4RZ1_B 4S1G_A 4S1G_B 4XX3_A 4XX3_B 4XX4_A 4XX4_B 5KOQ_A 5KOQ_B 5KOS_B 5KOT_B 5SXN_A 5SXN_B 5SY2_A 5SY2_B 5SY3_A 5SY3_B 5SZ9_A 5SZ9_B 5T4S_B 5TMG_A 5TMG_B 5TMK_B | 1BIL_A | 1BBS_A 1BBS_B 2BKS_A 2BKS_B 2BKT_A 2BKT_B 2FS4_A 2FS4_B 2G21_A 2G21_B 2G24_A 2G24_B 2G26_A 2G26_B 2G27_A 2G27_B 2V16_C 3G72_A 3GW5_B 3KM4_B 3VYF_A 5KOS_A 5KOT_A 5T4S_A 5TMK_A 5VPM_A 5VPM_B | 232 - 237 | 232 - 237 |
| P02699 | 1HZX_A 1L9H_A 1U19_A 2G87_A 2HPY_A 2PED_A | 1HZX_A | 1F88_A 1F88_B 1GZM_A 1GZM_B 1HZX_B 1L9H_B 3CAP_A 3CAP_B 3PQR_A 4J4Q_A 4PXF_A 4X1H_A | 327 - 348 | 327 - 348 |

Continues on next page.

| UniprotID | PdbId | PdbId (g(r) analysis) | PDB complex, not structured | Fuzzy Residues (Uniprot) | Fuzzy Residues (Pdb) |
|---|---|---|---|---|---|
| P02768 | 3A73_A 3LU6_A 3LU7_A 3LU7_B 3LU8_A 3LU8_B 4EMX_A 4EMX_B 5ID7_A | 3A73_A | 1TF0_A 2BXA_A 2BXA_B 2BXE_A 2BXE_B 2VDB_A 2VUE_A 2VUE_B 2XVQ_A 2XVQ_B 2XVU_A 2XVU_B 2XW0_A 2XW0_B 2XW1_A 2XW1_B 2YDF_A 2YDF_B 4HGM_B | 101 - 113 | 101 - 113 |
| P06730 | 2W97_A 3U7X_A 3U7X_B 4AZA_A | 2W97_A | 2V8W_A 2V8W_E 2V8X_A 2V8X_E 2V8Y_A 2V8Y_E 3TF2_A 3TF2_B 3TF2_C 3TF2_D 4AZA_C 4BEA_A 5EHC_A 5EIR_A | 206 - 210 | 206 - 210 |
| P07024 | 1HPU_B 1HPU_C 2USH_B | 1HPU_B | 2USH_A | 324 - 329 | 324 - 329 |
| P07607 | 4EB4_A 4EB4_B 4EB4_C 4EB4_D | 4EB4_A | 3IHI_A 3IHI_B 4E5O_A 4E5O_B 4E5O_C 4E5O_D 4E5O_E 4E5O_F 4EIN_A 4EIN_B | 1 - 13 | 1 - 13 |
| P08254 | 1G4K_A 1QIC_D 2D1O_A 2D1O_B | 1G4K_A | 1G05_A 1G05_B 1HY7_A 1HY7_B | 242 - 247 | 242 - 247 |
| P09391 | 5MT6_A 5MT7_A 5MT8_A | 5MT6_A | 2IRV_A 2NRF_B | 245 - 250 | 245 - 250 |
| P0A3F4 | 2V5H_G 2V5H_H 2V5H_I 2V5H_J 2V5H_K 2V5H_L | 2V5H_G | 4C3M_A 4C3M_B 4C3M_C | 37 - 53 | 37 - 53 |

Continues on next page.

| UniprotID | PdbId | PdbId (g(r) analysis) | PDB complex, not structured | Fuzzy Residues (Uniprot) | Fuzzy Residues (Pdb) |
|---|---|---|---|---|---|
| P0A6Z6 | 1Q5Y_A 1Q5Y_B 1Q5Y_C 1Q5Y_D 3BKT_A 3BKT_B 3BKT_C 3BKT_D 3BKU_A 3BKU_B 3BKU_C | 1Q5Y_A | 3BKU_D | 63 - 78 | 63 - 78 |
| P0C6U8 | 3SND_B 3VB3_B 3VB4_B 3VB5_B 3VB6_B 3VB7_B | 3SND_B | 3VB3_A 3VB4_A 3VB5_A 3VB6_A 3VB7_A | 3542 - 3546 | 3542 - 3546 |
| P12004 | 1UL1_B 1UL1_C 1VYJ_C 1VYJ_E 1VYJ_I 1VYJ_K 1VYM_B 4ZTD_A 4ZTD_B 5MLW_A 5MLW_E | 1U76_A | 1AXC_A 1AXC_C 1AXC_E 1U76_A 1U76_C 1U76_E 1U7B_A 1UL1_A 2ZVK_A 2ZVK_B 2ZVK_C 2ZVL_A 2ZVL_B 2ZVL_C 2ZVL_D 2ZVL_E 2ZVL_F 2ZVM_A 2ZVM_B 2ZVM_C 3P87_A 3P87_B 3P87_C 3P87_D 3P87_E 3P87_F 3TBL_A 3TBL_B 3TBL_C 3WGW_A 3WGW_B 4RJF_C 4RJF_E 5MLW_C | 256 - 260 | 256 - 260 |

Continues on next page.

| UniprotID | PdbId | PdbId (g(r) analysis) | PDB complex, not structured | Fuzzy Residues (Uniprot) | Fuzzy Residues (Pdb) |
|---|---|---|---|---|---|
| P12004 | 1UL1_B 1UL1_C 1VYJ_C 1VYJ_E 1VYJ_I 1VYJ_K 1VYM_B 4ZTD_A 4ZTD_B 5MLW_A 5MLW_E | 1UL1_B | 1AXC_A 1AXC_C 1AXC_E 1U76_A 1U76_C 1U76_E 1U7B_A 1UL1_A 2ZVK_A 2ZVK_B 2ZVK_C 2ZVL_A 2ZVL_B 2ZVL_C 2ZVL_D 2ZVL_E 2ZVL_F 2ZVM_A 2ZVM_B 2ZVM_C 3P87_A 3P87_B 3P87_C 3P87_D 3P87_E 3P87_F 3TBL_A 3TBL_B 3TBL_C 3WGW_A 3WGW_B 4RJF_C 4RJF_E 5MLW_C | 186 - 192 | 186 - 192 |
| P13298 | 2PS1_A 2PS1_B | 2PS1_A | 2PRZ_A 2PRZ_B 2PRZ_C 2PRZ_D | 110 - 115 | 110 - 115 |
| P13482 | 2JJB_A 2JJB_B 2JJB_D 2WYN_A 2WYN_B 2WYN_D | 2JJB_A | 2JJB_C 2WYN_C | 32 - 36 | 32 - 36 |
| P15369 | 2IFW_A 2IFW_B | 2IFW_A | 1S2K_A 2IFR_A | 130 - 134 | 130 - 134 |
| P18206 | 5L0D_A 5L0D_D | 5L0D_A | 5L0D_B 5L0D_C | 1117 - 1122 | 1117 - 1122 |
| P22894 | 1JAN_A | 1JAN_A | 1JAP_A | 100 - 104 | 100 - 104 |

Continues on next page.

| UniprotID | PdbId | PdbId (g(r) analysis) | PDB complex, not structured | Fuzzy Residues (Uniprot) | Fuzzy Residues (Pdb) |
|---|---|---|---|---|---|
| P24941 | 1F5Q_A 1F5Q_C 1FIN_A 1FIN_C 1FVV_A 1FVV_C 1OKV_A 1OKV_C 1OKW_A 1OKW_C 1OL1_A 1OL1_C 1OL2_A 1OL2_C 1URC_A 1URC_C 2C5N_A 2C5N_C 2C5O_A 2C5O_C 2C5V_A 2C5V_C 2C5X_A 2C5X_C 2I40_A 2I40_C 2UUE_A 2UUE_C 2V22_A 2V22_C 2WEV_A 2WEV_C 2WFY_A 2WFY_C 2WHB_A 2WHB_C 2X1N_A 2X1N_C 3F5X_A 3F5X_C 4FX3_A 4FX3_C 5IF1_A 5IF1_C | 1F5Q_A | 1BUH_A | 38 - 46 | 38 - 46 |

Continues on next page.

| UniprotID | PdbId | PdbId (g(r) analysis) | PDB complex, not structured | Fuzzy Residues (Uniprot) | Fuzzy Residues (Pdb) |
|---|---|---|---|---|---|
| P29476 | 4JSE_B | 4JSE_B | 1K2R_A 1K2R_B 1K2S_A 1K2S_B 1K2T_A 1K2T_B 1K2U_A 1K2U_B 1LZX_A 1LZX_B 1LZZ_A 1LZZ_B 1M00_A 1M00_B 1MMV_A 1MMV_B 1MMW_A 1MMW_B 1OM4_A 1OM4_B 1OM5_A 1OM5_B 1P6H_A 1P6H_B 1P6I_A 1P6I_B 1P6J_A 1P6J_B 1RS6_A 1RS6_B 1RS7_A 1RS7_B 2G6H_A 2G6H_B 2G6I_A 2G6I_B 2G6K_A 2G6K_B 2G6L_A 2G6L_B 2G6M_A 2G6M_B 2G6N_A 2G6N_B 2HX3_A 2HX3_B | 339 - 347 | 339 - 347 |

Continues on next page.

| UniprotID | PdbId | PdbId (g(r) analysis) | PDB complex, not structured | Fuzzy Residues (Uniprot) | Fuzzy Residues (Pdb) |
|---|---|---|---|---|---|
| P29476 | 4JSE_B | 4JSE_B | 2HX4_A 2HX4_B 3B3M_A 3B3M_B 3B3N_A 3B3N_B 3B3P_A 3B3P_B 3HSN_A 3HSN_B 3HSO_A 3HSO_B 3HSP_A 3HSP_B 3JT3_A 3JT3_B 3JT4_A 3JT4_B 3JT5_A 3JT5_B 3JT6_A 3JT6_B 3JT7_A 3JT7_B 3JT8_A 3JT8_B 3JT9_A 3JT9_B 3JTA_A 3JTA_B 3N2R_A 3N2R_B 3N5V_A 3N5V_B 3N5W_A 3N5W_B 3N5X_A 3N5X_B 3N5Y_A 3N5Y_B 3N5Z_A 3N5Z_B 3N60_A 3N60_B 3NLM_A 3NLM_B | 339 - 347 | 339 - 347 |

Continues on next page.

| UniprotID | PdbId | PdbId (g(r) analysis) | PDB complex, not structured | Fuzzy Residues (Uniprot) | Fuzzy Residues (Pdb) |
|---|---|---|---|---|---|
| P29476 | 4JSE_B | 4JSE_B | 3NLV_A<br>3NLV_B<br>3NLW_A<br>3NLW_B<br>3NLX_A<br>3NLX_B<br>3NLY_A<br>3NLY_B<br>3NLZ_A<br>3NLZ_B<br>3NM0_A<br>3NM0_B<br>3NNY_A<br>3NNY_B<br>3NNZ_A<br>3NNZ_B<br>3PNE_A<br>3PNE_B<br>3PNF_A<br>3PNF_B<br>3PNG_A<br>3PNG_B<br>3Q99_A 3Q99_B<br>3Q9A_A<br>3Q9A_B<br>3RQJ_A<br>3RQJ_B<br>3RQK_A<br>3RQK_B<br>3RQL_A<br>3RQL_B<br>3RQM_A<br>3RQM_B<br>3RQN_A<br>3RQN_B<br>3SVP_A<br>3SVP_B<br>3SVQ_A<br>3SVQ_B<br>3TYL_A<br>3TYL_B<br>3TYM_A<br>3TYM_B<br>3TYN_A<br>3TYN_B | 339 - 347 | 339 - 347 |

Continues on next page.

| UniprotID | PdbId | PdbId (g(r) analysis) | PDB complex, not structured | Fuzzy Residues (Uniprot) | Fuzzy Residues (Pdb) |
|---|---|---|---|---|---|
| P29476 | 4JSE_B | 4JSE_B | 3TYO_A<br>3TYO_B<br>3UFO_A<br>3UFO_B<br>3UFP_A<br>3UFP_B<br>3UFQ_A<br>3UFQ_B<br>3UFR_A<br>3UFR_B<br>3UFS_A<br>3UFS_B<br>3UFT_A<br>3UFT_B<br>3UFU_A<br>3UFU_B<br>3UFV_A<br>3UFV_B<br>3UFW_A<br>3UFW_B<br>4C39_A 4C39_B<br>4CAM_A<br>4CAM_B<br>4CAN_A<br>4CAN_B<br>4CAO_A<br>4CAO_B<br>4CAP_A<br>4CAP_B<br>4CAQ_A<br>4CAQ_B<br>4CDT_A<br>4CDT_B<br>4CTP_A<br>4CTP_B<br>4CTQ_A<br>4CTQ_B<br>4CTR_A<br>4CTR_B<br>4CTT_A<br>4CTT_B<br>4CTU_A<br>4CTU_B<br>4CTV_A<br>4CTV_B | 339 - 347 | 339 - 347 |

Continues on next page.

| UniprotID | PdbId | PdbId (g(r) analysis) | PDB complex, not structured | Fuzzy Residues (Uniprot) | Fuzzy Residues (Pdb) |
|---|---|---|---|---|---|
| P29476 | 4JSE_B | 4JSE_B | 4CTW_A 4CTW_B 4CTX_A 4CTX_B 4D2Y_A 4D2Y_B 4D2Z_A 4D2Z_B 4D30_A 4D30_B 4D31_A 4D31_B 4D32_A 4D32_B 4D3B_A 4D3B_B 4D7O_A 4D7O_B 4EUX_A 4EUX_B 4FVW_A 4FVW_B 4FVX_A 4FVX_B 4FVY_A 4FVY_B 4FVZ_A 4FVZ_B 4FW0_A 4FW0_B 4GQE_A 4GQE_B 4IMS_A 4IMS_B 4IMT_A 4IMT_B 4IMU_A 4IMU_B 4IMW_A 4IMW_B 4JSE_A 4JSF_A 4JSF_B 4JSI_A 4JSI_B 4JSJ_A 4JSJ_B | 339 - 347 | 339 - 347 |

Continues on next page.

| UniprotID | PdbId | PdbId (g(r) analysis) | PDB complex, not structured | Fuzzy Residues (Uniprot) | Fuzzy Residues (Pdb) |
|---|---|---|---|---|---|
| P29476 | 4JSE_B | 4JSE_B | 4K5D_A 4K5D_B 4K5F_A 4K5F_B 4K5G_A 4K5G_B 4KCH_A 4KCH_B 4KCI_A 4KCI_B 4KCJ_A 4KCJ_B 4KCK_A 4KCK_B 4KCL_A 4KCL_B 4KCM_A 4KCM_B 4KCN_A 4KCN_B 4KCO_A 4KCO_B 4LUX_A 4LUX_B 4UGZ_A 4UGZ_B 4UH0_A 4UH0_B 4UH1_A 4UH1_B 4UH2_A 4UH2_B 4UH3_A 4UH3_B 4UH4_A 4UH4_B 4UPM_A 4UPM_B 4UPN_A 4UPN_B 4UPO_A 4UPO_B 4UPP_A 4UPP_B 4V3V_A 4V3V_B | 339 - 347 | 339 - 347 |

Continues on next page.

| UniprotID | PdbId | PdbId (g(r) analysis) | PDB complex, not structured | Fuzzy Residues (Uniprot) | Fuzzy Residues (Pdb) |
|---|---|---|---|---|---|
| P29476 | 4JSE_B | 4JSE_B | 4V3W_A 4V3W_B 4V3X_A 4V3X_B 4V3Y_A 4V3Y_B 4V3Z_A 4V3Z_B 5AD4_A 5AD4_B 5AD5_A 5AD5_B 5AD6_A 5AD6_B 5AD7_A 5AD7_B 5AD8_A 5AD8_B 5AD9_A 5AD9_B 5ADA_A 5ADA_B 5ADB_A 5ADB_B 5ADC_A 5ADC_B 5AGK_A 5AGK_B 5AGL_A 5AGL_B 5AGM_A 5AGM_B 5AGN_A 5AGN_B 5AGO_A 5AGO_B 5AGP_A 5AGP_B 5FVP_A 5FVP_B 5FVQ_A 5FVQ_B 5FVR_A 5FVR_B 5FVS_A 5FVS_B | 339 - 347 | 339 - 347 |

Continues on next page.

| UniprotID | PdbId | PdbId (g(r) analysis) | PDB complex, not structured | Fuzzy Residues (Uniprot) | Fuzzy Residues (Pdb) |
|---|---|---|---|---|---|
| P29476 | 4JSE_B | 4JSE_B | 5FVT_A<br>5FVT_B<br>5FW0_A<br>5FW0_B<br>5UNR_A<br>5UNR_B<br>5UNS_A<br>5UNS_B<br>5UNT_A<br>5UNT_B<br>5UNU_A<br>5UNU_B<br>5UNV_A<br>5UNV_B<br>5UNW_A<br>5UNW_B<br>5UNX_A<br>5UNX_B<br>5UNY_A<br>5UNY_B<br>5UNZ_A<br>5UNZ_B<br>5UO0_A<br>5UO0_B<br>5VUI_A 5VUI_B<br>5VUJ_A<br>5VUJ_B<br>5VUK_A<br>5VUK_B<br>5VUL_A<br>5VUL_B<br>5VUM_A<br>5VUM_B<br>5VUN_A<br>5VUN_B<br>5VUO_A<br>5VUO_B<br>5VUP_A<br>5VUP_B<br>5VUQ_A<br>5VUQ_B<br>5VUR_A<br>5VUR_B<br>5VUS_A<br>5VUS_B<br>5VUT_A<br>5VUT_B<br>5VUU_A<br>5VUU_B | 339 - 347 | 339 - 347 |

Continues on next page.

| UniprotID | PdbId | PdbId (g(r) analysis) | PDB complex, not structured | Fuzzy Residues (Uniprot) | Fuzzy Residues (Pdb) |
|---|---|---|---|---|---|
| P29563 | 1VGO_B 2Z55_A 2Z55_D | 1VGO_B | 1VGO_A 2Z55_B 2Z55_E | 244 - 248 | 244 - 248 |
| P31153 | 4NDN_B 4NDN_D | 4NDN_B | 4NDN_A 4NDN_C | 116 - 126 | 116 - 126 |
| P31570 | 1G64_B | 1G64_B | 1G64_A | 7 - 26 | 7 - 26 |
| P33284 | 3O08_B 3O1W_A 3O1W_B 3O4W_A 3O4W_B 3O5B_A 3O5B_B | 3O08_B | 3O08_A 3O6W_A 3O6W_B | 2 - 16 | 2 - 16 |
| P37231 | 2ATH_B 2F4B_B 2G0G_B 2HWQ_B 2HWR_B 3GBK_B 3NOA_B | 2ATH_B | 1KNU_A 1KNU_B 1NYX_A 1NYX_B 2FVJ_A 2GTK_A 2HWQ_A 2HWR_A 2POB_A 2POB_B 2Q8S_A 2Q8S_B 3B0Q_A 3B0Q_B 3B0R_A 3B0R_B 3CWD_A 3CWD_B 3FEJ_A 3FUR_A 3G9E_A 3H0A_D 3IA6_A 3IA6_B 3K8S_A 3K8S_B 3KMG_A 3KMG_D 3U9Q_A 4FGY_A 4R06_A 4R06_B 4R2U_A 4R2U_D 4Y29_A 5GTN_A 5GTO_A 5GTP_A 5TTO_A 5TTO_B 5U5L_A 5U5L_B | 293 - 301 | 293 - 301 |
| P38424 | 1SVW_B | 1SVW_B | 1SUL_A | 51 - 59 | 51 - 59 |
| P42336 | 4L1B_A 4L23_A 4L2Y_A | 4L1B_A | 5DXH_A 5DXH_D | 414 - 418 | 414 - 418 |

Continues on next page.

| UniprotID | PdbId | PdbId (g(r) analysis) | PDB complex, not structured | Fuzzy Residues (Uniprot) | Fuzzy Residues (Pdb) |
|---|---|---|---|---|---|
| P42336 | 4L1B_A 4L23_A 4L2Y_A | 4L1B_A | 5DXH_A 5DXH_D | 940 - 948 | 940 - 948 |
| P42700 | 1YVP_A 1YVP_B | 1YVP_A | 2I91_A 2I91_B | 136 - 143 | 136 - 143 |
| P43912 | 4MCB_B | 4MCB_B | 4MCB_A 4MCC_A 4MCC_B | 161 - 169 | 161 - 169 |
| P49137 | 2PZY_C 2PZY_D 3A2C_A 3A2C_B 3A2C_C 3A2C_D 3A2C_E 3A2C_F 3A2C_H 3A2C_I 3A2C_J 3A2C_K 3A2C_L 3KC3_A 3KC3_D 3KC3_E 3KC3_F 3KC3_H 3KC3_I 3KC3_K 3KC3_L 4TYH_A | 2PZY_C | 2PZY_B 3A2C_G 3KC3_B 3KC3_C 3KC3_G 3KC3_J 3R2B_A 3R2B_B 3R2B_C 3R2B_D 3R2B_E 3R2B_F 3R2B_G 3R2B_H 3R2B_I 3R2B_J 3R2B_K 3R2B_L 3WI6_A 3WI6_B 3WI6_C 3WI6_D 3WI6_E 3WI6_F | 265 - 274 | 265 - 274 |
| P49137 | 2PZY_C 2PZY_D 3A2C_A 3A2C_B 3A2C_C 3A2C_D 3A2C_E 3A2C_F 3A2C_H 3A2C_I 3A2C_J 3A2C_K 3A2C_L 3KC3_A 3KC3_D 3KC3_E 3KC3_F 3KC3_H 3KC3_I 3KC3_K 3KC3_L 4TYH_A | 2PZY_C | 2PZY_B 3A2C_G 3KC3_B 3KC3_C 3KC3_G 3KC3_J 3R2B_A 3R2B_B 3R2B_C 3R2B_D 3R2B_E 3R2B_F 3R2B_G 3R2B_H 3R2B_I 3R2B_J 3R2B_K 3R2B_L 3WI6_A 3WI6_B 3WI6_C 3WI6_D 3WI6_E 3WI6_F | 346 - 364 | 346 - 364 |

Continues on next page.

| UniprotID | PdbId | PdbId (g(r) analysis) | PDB complex, not structured | Fuzzy Residues (Uniprot) | Fuzzy Residues (Pdb) |
|---|---|---|---|---|---|
| P49137 | 2PZY_C 2PZY_D 3A2C_A 3A2C_B 3A2C_C 3A2C_D 3A2C_E 3A2C_F 3A2C_H 3A2C_I 3A2C_J 3A2C_K 3A2C_L 3KC3_A 3KC3_D 3KC3_E 3KC3_F 3KC3_H 3KC3_I 3KC3_K 3KC3_L 4TYH_A | 3R2B_C | 2PZY_B 3A2C_G 3KC3_B 3KC3_C 3KC3_G 3KC3_J 3R2B_A 3R2B_B 3R2B_C 3R2B_D 3R2B_E 3R2B_F 3R2B_G 3R2B_H 3R2B_I 3R2B_J 3R2B_K 3R2B_L 3WI6_A 3WI6_B 3WI6_C 3WI6_D 3WI6_E 3WI6_F | 153 - 158 | 153 - 158 |
| P49137 | 2PZY_C 3KC3_B 3KC3_F 3KC3_H 3KC3_I 4TYH_A | 2PZY_A | 2PZY_A 2PZY_B 2PZY_D 3A2C_A 3A2C_B 3A2C_C 3A2C_D 3A2C_E 3A2C_F 3A2C_G 3A2C_H 3A2C_I 3A2C_J 3A2C_K 3A2C_L 3KC3_A 3KC3_C 3KC3_D 3KC3_E 3KC3_G 3KC3_J 3KC3_K 3KC3_L 3R2B_A 3R2B_B 3R2B_C 3R2B_D 3R2B_E 3R2B_F 3R2B_G 3R2B_H 3R2B_I 3R2B_J 3R2B_K 3R2B_L 3WI6_A 3WI6_B 3WI6_C 3WI6_D 3WI6_E 3WI6_F | 216 - 229 | 216 - 229 |

Continues on next page.

| UniprotID | PdbId | PdbId (g(r) analysis) | PDB complex, not structured | Fuzzy Residues (Uniprot) | Fuzzy Residues (Pdb) |
|---|---|---|---|---|---|
| P49137 | 2PZY_C 3KC3_B 3KC3_F 3KC3_H 3KC3_I 4TYH_A | 2PZY_A | 2PZY_A 2PZY_B 2PZY_D 3A2C_A 3A2C_B 3A2C_C 3A2C_D 3A2C_E 3A2C_F 3A2C_G 3A2C_H 3A2C_I 3A2C_J 3A2C_K 3A2C_L 3KC3_A 3KC3_C 3KC3_D 3KC3_E 3KC3_G 3KC3_J 3KC3_K 3KC3_L 3R2B_A 3R2B_B 3R2B_C 3R2B_D 3R2B_E 3R2B_F 3R2B_G 3R2B_H 3R2B_I 3R2B_J 3R2B_K 3R2B_L 3WI6_A 3WI6_B 3WI6_C 3WI6_D 3WI6_E 3WI6_F | 236 - 240 | 236 - 240 |

Continues on next page.

| UniprotID | PdbId | PdbId (g(r) analysis) | PDB complex, not structured | Fuzzy Residues (Uniprot) | Fuzzy Residues (Pdb) |
|---|---|---|---|---|---|
| P52732 | 2FL2_A 2FL6_A 2Q2Z_A 4A5Y_C 4BXN_A | 2FL2_A | 1II6_A 1II6_B 1Q0B_A 1Q0B_B 1X88_A 1X88_B 1YRS_A 1YRS_B 2FKY_A 2FKY_B 2FL2_B 2FL6_B 2FME_A 2FME_B 2G1Q_A 2G1Q_B 2GM1_A 2GM1_B 2GM1_D 2GM1_E 2IEH_A 2IEH_B 2PG2_A 2PG2_B 2Q2Y_A 2Q2Y_B 2Q2Z_B 2UYI_A 2UYI_B 2UYM_A 2UYM_B 2WOG_A 2WOG_B 2WOG_C 2X2R_A 2X2R_B 2X2R_C 2X7C_A 2X7C_B 2X7D_A 2X7D_B 2X7E_A 2X7E_B 2XAE_A 2XAE_B 2XAE_C 3CJO_A 3CJO_B 3K3B_A 3K3B_B 3K5E_A 3K5E_B 3L9H_A 3L9H_B 4A51_A 4A51_B 4A51_C 4A51_D 4A51_E 4A51_F 4A51_G 4A5Y_A 4A5Y_B | 272 - 287 | 272 - 287 |

Continues on next page.

| UniprotID | PdbId | PdbId (g(r) analysis) | PDB complex, not structured | Fuzzy Residues (Uniprot) | Fuzzy Residues (Pdb) |
|---|---|---|---|---|---|
| P53350 | 1Q4K_C 3HIH_A 4HAB_A | 1Q4K_C | 1Q4O_A 1Q4O_B 2OJX_A 3C5L_A 3HIH_B 4HAB_C 4HY2_A 4LKL_A 4LKM_A 4LKM_C 4O9W_A 4RCP_A | 495 - 506 | 495 - 506 |
| P53779 | 3V6R_A 3V6S_A 4H3B_A | 3V6R_A | 3OXI_A 3V6R_B 3V6S_B | 211 - 224 | 211 - 224 |
| P53779 | 3V6R_A 3V6S_A 4H3B_A | 3V6R_A | 3OXI_A 3V6R_B 3V6S_B | 369 - 382 | 369 - 382 |
| P56817 | 1M4H_A 1SGZ_C 1SGZ_D 1XN2_A 1XN3_B 1XN3_C 5V0N_B 5V0N_C | 1M4H_A | 3IXJ_A 3IXJ_B 3IXJ_C 3IXK_A 3IXK_B 3IXK_C 4K8S_A 4K8S_B 4K8S_C 4K9H_A 4K9H_B 4K9H_C 5DQC_A 5DQC_B 5DQC_C 5V0N_A | 218 - 230 | 218 - 230 |

Continues on next page.

| UniprotID | PdbId | PdbId (g(r) analysis) | PDB complex, not structured | Fuzzy Residues (Uniprot) | Fuzzy Residues (Pdb) |
|---|---|---|---|---|---|
| P61823 | 1A2W_A 1A2W_B 1AFK_B 1AFL_B 1AFU_B 1DFJ_E 1EOS_B 1JN4_B 1JVT_B 1JVU_B 1JVV_B 1O0F_B 1O0H_B 1O0M_B 1O0N_B 1O0O_B 1QHC_B 1RBB_A 1U1B_B 1W4O_B 1W4P_B 1W4Q_B 1XPS_A 1XPT_A 1Z6D_B 1Z6S_B 2G4W_B 2G8Q_B 2G8R_B 2P42_A 2P42_C 2W5G_B 2W5I_B 2W5K_B 2W5L_B 2W5M_B 2XOG_B 3D6O_B 3D6P_B 3D6Q_B 3D7B_B 3D8Y_B 3D8Z_B 3DXG_B 3DXH_B 3EUX_B 3EUY_B 3EUZ_B 3EV0_B 3EV1_B 3EV2_B 3EV3_B 3EV4_B 3EV5_B 3EV6_B 3JW1_B 4G8V_B 4G8Y_B 4G90_B 4L55_B 4MXF_B 4OT4_A 4PEQ_A 4PEQ_C 4QH3_B 4S0Q_B 5E5E_B 5E5F_B 5JLG_B 5JMG_B 5JML_B 5OBC_B 5OBD_B 5OBE_B | 1A2W_A | 2P44_A 2P49_A 2XOG_A 2XOI_B 3EUX_A 3EUY_A 3EUZ_A 3EV1_A 3EV2_A 3EV3_A 3EV4_A 3EV6_A 3QSK_A 4OT4_B 4POU_A 4QH3_A 4S0Q_A 4S18_A 4S18_B | 42 - 49 | 42 - 49 |

Continues on next page.

| UniprotID | PdbId | PdbId (g(r) analysis) | PDB complex, not structured | Fuzzy Residues (Uniprot) | Fuzzy Residues (Pdb) |
|---|---|---|---|---|---|
| P61964 | 4GM9_A 4GM9_B | 4GM9_A | 2H68_A 2H68_B 2H6K_A 2H6K_B 2H6N_A 2H6N_B 2H6Q_A 2H6Q_B 3EG6_A 4ERQ_A 4ERQ_B 4ERQ_C 4ERY_A 4ERZ_A 4ERZ_B 4ERZ_C 4ES0_A 4ESG_A 4ESG_B 4EWR_A 4GM8_A 4GM8_B 4GM8_C 4GM8_D 4GMB_A 4Y7R_A | 24 - 30 | 24 - 30 |

Continues on next page.

| UniprotID | PdbId | PdbId (g(r) analysis) | PDB complex, not structured | Fuzzy Residues (Uniprot) | Fuzzy Residues (Pdb) |
|---|---|---|---|---|---|
| P68135 | 1IJJ_B 1RGI_A 2A40_A 2A40_D 2FF6_A 2Q1N_B 3FFK_B 3FFK_E 4B1V_B 4K42_A 4K42_B 4PKG_A 4PKH_A 4PKH_D 4PKH_F 4PKI_A 4PL8_A 4Z94_A | 1IJJ_B | 1IJJ_A 1KXP_A 1LOT_B 1P8Z_A 1RFQ_A 1RFQ_B 1SQK_A 2FF3_B 2Q1N_A 2Q31_A 2Q31_B 2Q97_A 2V51_B 2V51_D 2V52_B 2VYP_A 2VYP_B 3BUZ_B 3M1F_A 3M6G_A 3M6G_B 3SJH_A 3U8X_A 3U8X_C 3U9Z_A 3UE5_A 4B1V_A 4B1X_B 4B1Y_B 4GY2_B 4H03_B 4H0T_B 4H0V_B 4H0X_B 4H0Y_B 4K42_C 4K42_D 4K43_A 4K43_B 4PKH_I 4PL8_B | 40 - 67 | 40 - 67 |

Continues on next page.

| UniprotID | PdbId | PdbId (g(r) analysis) | PDB complex, not structured | Fuzzy Residues (Uniprot) | Fuzzy Residues (Pdb) |
|---|---|---|---|---|---|
| P68135 | 1IJJ_B 1RGI_A 2A40_A 2A40_D 2FF6_A 2Q1N_B 3FFK_B 3FFK_E 4B1V_B 4K42_A 4K42_B 4PKG_A 4PKH_A 4PKH_D 4PKH_F 4PKI_A 4PL8_A 4Z94_A | 3M1F_A | 1IJJ_A 1KXP_A 1LOT_B 1P8Z_A 1RFQ_A 1RFQ_B 1SQK_A 2FF3_B 2Q1N_A 2Q31_A 2Q31_B 2Q97_A 2V51_B 2V51_D 2V52_B 2VYP_A 2VYP_B 3BUZ_B 3M1F_A 3M6G_A 3M6G_B 3SJH_A 3U8X_A 3U8X_C 3U9Z_A 3UE5_A 4B1V_A 4B1X_B 4B1Y_B 4GY2_B 4H03_B 4H0T_B 4H0V_B 4H0X_B 4H0Y_B 4K42_C 4K42_D 4K43_A 4K43_B 4PKH_I 4PL8_B | 3 - 7 | 3 - 7 |
| P70731 | 3MHY_A 3MHY_B 3MHY_C 4CNZ_A 4CNZ_C 4CNZ_D 4CNZ_E 4CNZ_F 4CO0_A 4CO0_B | 3MHY_A | 3O5T_B 4CNZ_B 4CO3_B 4CO4_A 4CO4_B 4CO4_C | 38 - 53 | 38 - 53 |
| P71707 | 5CRF_A 5CRF_B 5CRF_C | 5CRF_A | 5CRF_D | 775 - 779 | 775 - 779 |
| P98170 | 2OPZ_A 2OPZ_B 2OPZ_C 2OPZ_D | 2OPZ_A | 1G73_C | 347 - 356 | 347 - 356 |
| P9WPN8 | 5LIE_B | 5LIE_B | 5LI6_A 5LI6_B 5LI7_A 5LI7_B 5LIE_A | 179 - 192 | 179 - 192 |
| Q02248 | 1I7W_A 1I7W_C 1I7X_C | 1I7W_A | 1I7X_A 1M1E_A 1V18_A 4EVA_A 4EVA_C | 134 - 161 | 134 - 161 |
| Q02248 | 1I7W_A 1I7W_C 1I7X_C | 1M1E_A | 1I7X_A 1M1E_A 1V18_A 4EVA_A 4EVA_C | 663 - 671 | 663 - 671 |

Continues on next page.

| UniprotID | PdbId | PdbId (g(r) analysis) | PDB complex, not structured | Fuzzy Residues (Uniprot) | Fuzzy Residues (Pdb) |
|---|---|---|---|---|---|
| Q06187 | 1K2P_A 1K2P_B 3P08_B 5J87_A 5J87_C | 1K2P_A | 3P08_A 5FBN_C 5FBN_D | 545 - 556 | 545 - 556 |
| Q06755 | 5DDT_A 5DDT_B | 5DDT_A | 5HS2_A 5HS2_B | 227 - 232 | 227 - 232 |
| Q09028 | 4PBY_B 4PC0_A 4PC0_B | 2XU7_B | 2XU7_A 2XU7_B 4PBY_A 4PBZ_A 4R7A_B | 55 - 59 | 55 - 59 |
| Q09028 | 4PBY_B 4PC0_A 4PC0_B | 4PBY_A | 2XU7_A 2XU7_B 4PBY_A 4PBZ_A 4R7A_B | 2 - 15 | 2 - 15 |
| Q09028 | 4PBY_B 4PC0_A 4PC0_B | 4PBY_A | 2XU7_A 2XU7_B 4PBY_A 4PBZ_A 4R7A_B | 353 - 360 | 353 - 360 |
| Q09028 | 4PBY_B 4PC0_A 4PC0_B | 4PBY_B | 2XU7_A 2XU7_B 4PBY_A 4PBZ_A 4R7A_B | 89 - 113 | 89 - 113 |
| Q14012 | 4FG8_B | 4FG8_B | 4FG8_A 4FG9_A 4FG9_B | 172 - 179 | 172 - 179 |
| Q41460 | 3CNZ_A 3CNZ_B | 3CNZ_A | 3COB_A 3COB_C | 1234 - 1240 | 1234 - 1240 |
| Q46893 | 3N9W_B | 3N9W_B | 3N9W_A | 229 - 235 | 229 - 235 |
| Q4U1A6 | 3AGD_A 3AGD_B 3AGE_A 3AGE_B | 3AGD_A | 3IH8_A 3IH8_B 3IHA_A 3IHA_B | 354 - 380 | 354 - 380 |
| Q57872 | 1PKH_A 1PKJ_A 1PKK_A | 1PKH_A | 1OGH_A 1OGH_B 1PKH_B 1PKJ_B 1PKK_B | 176 - 182 | 176 - 182 |
| Q5JII7 | 3VYS_C 3VYT_C | 3VYS_C | 3VYU_C | 15 - 19 | 15 - 19 |
| Q5JII7 | 3VYS_C 3VYT_C | 3VYS_C | 3VYU_C | 3 - 10 | 3 - 10 |
| Q6ST21 | 2EPN_A 2EPN_B 2EPO_A | 2EPN_A | 2EPO_B | 273 - 287 | 273 - 287 |
| Q7KQK5 | 2Z8V_A 2Z8W_A 4R19_B | 2Z8V_A | 4R19_A 4R1C_A 4R1C_B | 172 - 176 | 172 - 176 |
| Q7KQK5 | 2Z8V_A 2Z8W_A 4R19_B | 2Z8V_A | 4R19_A 4R1C_A 4R1C_B | 259 - 272 | 259 - 272 |
| Q7KQK5 | 2Z8V_A 2Z8W_A 4R19_B | 2Z8W_A | 4R19_A 4R1C_A 4R1C_B | 352 - 356 | 352 - 356 |
| Q7KQK5 | 2Z8V_A 2Z8W_A 4R19_B | 2Z8W_A | 4R19_A 4R1C_A 4R1C_B | 370 - 387 | 370 - 387 |

Continues on next page.

| UniprotID | PdbId | PdbId (g(r) analysis) | PDB complex, not structured | Fuzzy Residues (Uniprot) | Fuzzy Residues (Pdb) |
|---|---|---|---|---|---|
| Q84AF2 | 2P0J_A | 2P0J_A | 2P0J_B | 47 - 52 | 47 - 52 |
| Q84B82 | 3TCJ_B | 3TCJ_B | 4ELY_C 4ELY_D | 46 - 50 | 46 - 50 |
| Q84BQ9 | 3CJQ_A 3CJQ_D 3CJQ_G 3CJR_A 3EGV_A | 3CJQ_A | 2NXJ_A 2NXJ_B 2NXN_A | 97 - 101 | 97 - 101 |
| Q89ZI2 | 4UR9_A 4UR9_B 5FL0_A 5FL0_B 5FL1_A 5FL1_B | 4UR9_A | 2J4G_A 2J4G_B 2VVN_A 2VVN_B 2W66_A 2W66_B 2W67_A 2W67_B 2X0H_A 2X0H_B 2XJ7_A 2XJ7_B 2XM1_A 2XM1_B 2XM2_A 2XM2_B 4AIS_A 4AIS_B 5FKY_A 5FKY_B | 611 - 670 | 611 - 670 |
| Q8EMJ9 | 3ES7_A 3ES7_B 3ES8_A 3ES8_B 3ES8_C 3ES8_D 3ES8_E 3ES8_F 3ES8_G 3ES8_H | 3ES7_A | 2OQY_A 2OQY_B 2OQY_C 2OQY_D 2OQY_E 2OQY_F 2OQY_G 2OQY_H 3FYY_A 3FYY_B | 375 - 387 | 375 - 387 |
| Q93DA8 | 3EXS_D | 3EXS_D | 3EXR_B 3EXR_C 3EXS_A 3EXS_B 3EXS_C | 145 - 151 | 145 - 151 |
| Q97UA0 | 3BQB_X | 3BQB_X | 3BQB_A 3BQB_Y 3BQB_Z | 90 - 98 | 90 - 98 |
| Q980A5 | 3CW2_A 3CW2_E 3CW2_F 3I1F_A 4M0L_A | 3CW2_A | 2AHO_A 2QN6_A 3CW2_B 4M0L_B 4M0L_C 4M0L_D 4M0L_E 5DSZ_B | 39 - 47 | 39 - 47 |

Continues on next page.

| UniprotID | PdbId | PdbId (g(r) analysis) | PDB complex, not structured | Fuzzy Residues (Uniprot) | Fuzzy Residues (Pdb) |
|---|---|---|---|---|---|
| Q9BX68 | 4NJZ_B 4NJZ_C | 4NJZ_B | 4NK0_A 4NK0_B | 51 - 62 | 51 - 62 |
| Q9R194 | 4U8H_A | 4U8H_A | 4I6G_A 4I6G_B 4I6J_A | 248 - 257 | 248 - 257 |
| Q9U1E1 | 3NGS_C 3NGT_A 3NGT_B 3NGT_C 3NGT_D 3NGT_E 3NGT_F 3NGT_G 3NGT_H 3NGT_I 3NGT_J 3NGT_K 3NGT_N | 3NGS_C | 3NGT_L | 43 - 66 | 43 - 66 |

**Table 2.** Structures of complexes, which were generated by conditional folding of protein regions (PDB$_C DR$). PDB IDs for all complexes are listed, those used for g(r) analysis are shown in a separate column. The sequence numbers of DOR regions are given based on both Uniprot and PDB numbering.
